# Environmental DNA as an Indicator of Seasonal Reproductive Phenology in Freshwater Mussels

**DOI:** 10.64898/2026.02.19.706874

**Authors:** Nathaniel Marshall, Cheryl Dean, Mae Sierra, W. Cody Fleece

## Abstract

Unionid freshwater mussels exhibit a unique form of mitochondrial inheritance, termed doubly uniparental inheritance, in which a maternal and a paternal mitotype is transmitted uniparentally. The exclusive presence of a male mitotype in gonadal tissue and sperm cells suggests that environmental DNA (eDNA) could serve as a non-invasive method for monitoring freshwater mussel reproduction. Yet, the dynamics of male mitotype detection within the environment remain poorly understood. This study analyzed seasonal eDNA samples from two diverse mussel beds, detecting 24 mitochondrial operational taxonomic units (MOTUs) associated with the male mitotype. Peaks in male mitotype signal for mussels identifiable to the species level generally aligned with expected spawning periods based on female gravidity records (e.g., *Pyganodon grandis*, *Lasmigona costata*, *Ortmaniana ligamentina*). Additionally, male mitotype detection was often sporadic compared to the consistently detected female mitotype, indicating that male signals may be tied to behavioral or reproductive events rather than continuous shedding. While elevated male signals may reflect spawning, alternative sources such as tissue decay, mitotype leakage, glochidia release, or post-spawning gamete clearance complicate interpretation. A male-to-female mitotype ratio is proposed as a more reliable proxy for identifying sperm release events, given the high concentration of male mitotypes that occurs within spermatozeugmata. Limitations in male mitotype reference databases hindered species-level resolution for many MOTUs, underscoring the need for expanded genomic resources. Overall, this work demonstrates that male mitotype eDNA likely provides valuable insights into mussel reproductive ecology, while emphasizing the importance of long-term monitoring and integrated gametogenesis studies to refine its application in conservation.

## Introduction

Freshwater bivalves have faced massive declines globally, with freshwater mussels (Bivalvia: Unionidae) considered the most imperiled group of bivalves (Lopes-Lima et al. 2018, Aldridge et al. 2022). North America is comprised of the greatest freshwater mussel richness; however, >70% of these species are protected and deemed endangered, threatened, or species of concern (Williams et al. 2017, FMCS 2021). Habitat protection, restoration, and propagation programs have been developed to aid in conservation efforts for freshwater mussels (FMCS 2016, McMurray & Roe 2017), however monitoring these efforts is challenging (Strayer et al. 2019).

Freshwater mussels reproduce through spermcasting, where males discharge spermatozeugmata sperm masses consisting of thousands of spermatozoa (Waller & Lasee 1997, Ishibashi et al. 2000). Females then capture the spermatozeugmata from the water column and brood their glochidia larvae until they mature (Bishop & Pemberton 2006). The timing of spawning and brooding can vary by species and has previously been broadly categorized as either short-term (tachytictic) or long-term (bradytictic) (Haag 2012). Short-term brooders typically spawn in the late winter and early spring, with females brooding the larvae for a short period (2–8 weeks); whereas long-term brooders typically spawn in the late summer and autumn, with females brooding through the winter (Haag 2012). Watters and O’Dee (2000), after studying glochidial release, found the brooding categories were too simplistic to adequately encompass the behaviors observed in nature. While the phenology of spermcasting and fertilization has been studied (see Jirka & Neves 1992, Mosley et. al 2014, Wacker et al. 2018) timing is most often inferred from the inspection of gravid females.

In contrast to mitochondria (mt) strict maternal inheritance common across metazoans, freshwater mussels exhibit doubly uniparental inheritance (DUI), in which a maternal and a paternal mitochondrial (mt) genome is transmitted uniparentally (Zouros 2013, Soroka & Burzyński 2010, Gusman et al. 2016). The female mitotype occurs within female gonads and somatic tissues of both sexes, whereas the male mitotype typically occurs in high copy numbers within male gametes and gonads (with some exceptions, see Breton et al. 2017). The female and male mitotypes are easily distinguishable as the two mitotypes have greater than 40% intraspecific divergence (Doucet-Beaupré et al. 2010, Guerra et al. 2017).

Environmental DNA (eDNA – traces of shed genetic material found within the environment) has previously been useful in deciphering spawning events for fishes (Erickson et al. 2016, Tillotson et al. 2018, Thalinger et al. 2019) and a marine bivalve (Bayer et al. 2019). In the case of freshwater mussels, the DUI system provides a unique opportunity to track eDNA signals associated with male gametes (see Marshall et al. 2025). The concentration of the male mitotype is expected to be much higher within spermatozeugmata compared to other somatic tissue because (1) spermatozoa only consist of the male mitotype (Zouros 2013), (2) each individual spermatozoa contain several mt genomes (Waller and Lasee 1997), and (3) each spermatozeugmata consists of thousands of spermatozoa (Waller and Lasee 1997). Therefore, the presence of spermatozeugmata within the water column may be evident based on an increase in the presence of eDNA from the male mitotype.

Investigating spawning behaviors of freshwater mussels in natural environments presents significant challenges due to their cryptic habits, patchy distributions, and difficulties associated with field surveys. A rigorous understanding of reproductive biology requires reliable, non-lethal methods to determine sex, detect gametes, and confirm female gravidity, thereby minimizing impacts on vulnerable populations (Dudding et al. 2020). Therefore, the detection of male mitotype eDNA presents an opportunity for a non-invasive sampling methodology for understanding timing and environmental triggers of sperm release (Marshall et al. 2025). However, while freshwater mussel male mitotypes are known to be detectable within the environment, it is currently unknown how this detection varies seasonally and if these detections correlate with expected spawning behaviors. This study analyzed eDNA samples collected from two sites in central Ohio from April thru October, with a particular focus on detection of the male mitotype. This study assessed whether male mitotype signal fluctuated seasonally, if these patterns aligned with known spawning behaviors, and how male signal corresponded to detection trends of the female mitotype.

## Methods

### eDNA Sample Collection

Two sites in central Ohio, USA, were chosen to survey across a reproductive season for freshwater mussels (Table 1). The two sites were chosen based on known presence of high mussel diversity and the presence of multiple federally protected species. A total of ten sampling events occurred at the Killbuck Creek site, and nine total sampling events occurred at the Walhonding River site. The initial sampling event was on April 25, 2024 in Killbuck Creek and May 09, 2024 in Walhonding River. The initial sampling date was delayed in Walhonding River as river stage was too high for sampling in April. At both sites, the final sampling date was October 22, 2024 (Supplementary Table 1).

**Table 1.** Survey site information for eDNA sampling conducted in Killbuck Creek and Walhonding River.

| **Sampling Site** | **Latitude** | **Longitude** | **County** | **Drainage**<br>**(sq. miles)** | **USGS**<br>**Gage** | **Site Distance to**<br>**Gage** |
| --- | --- | --- | --- | --- | --- | --- |
| Killbuck Cr. | 40.40056 | -81.9575 | Holmes | 582 | 03139000 | 19 KM downstream |
| Walhonding R. | 40.32880 | -81.9896 | Coshocton | 1570 | 03139850 | 9 KM upstream |

On each sampling date, three replicate water samples were collected at Killbuck Creek and Walhonding River, with the exception of the August 29, 2024 sampling at Walhonding River which consisted of nine total water replicates (Supplementary Table 1). These nine water replicates were collected along the entire perimeter of the central island (Figure 1). The 10 sampling events in Killbuck Creek resulted in 30 total samples, while the nine sampling events in the Walhonding River resulted in 33 total samples (Supplementary Table 1).

**Figure 1.**
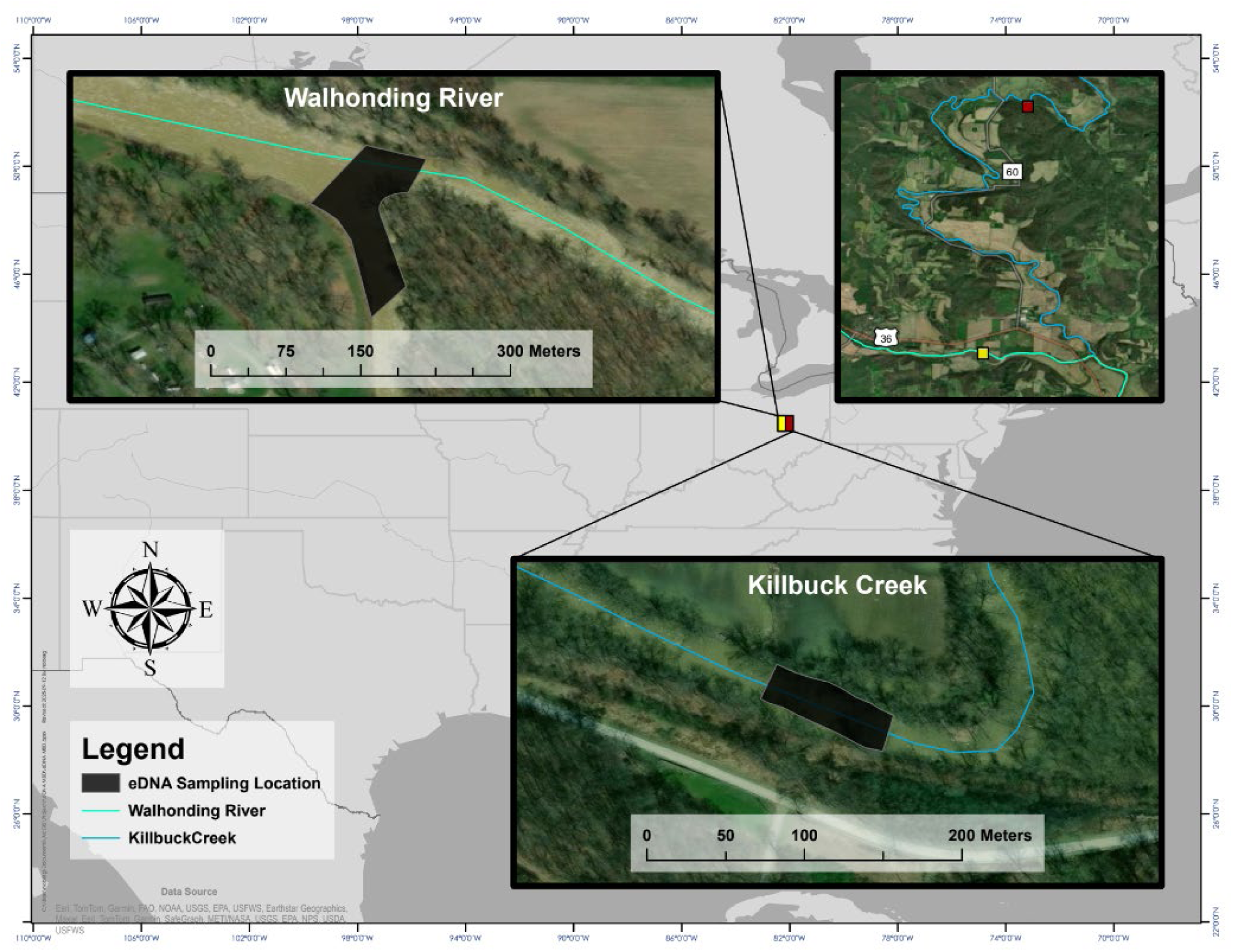
Map of eDNA survey sites sampled in the Walhonding River and Killbuck Creek.

Water samples were collected from near the benthos using a peristaltic pump with a polypropylene filter holder attached to a painter’s pole. A new polypropylene filter holder was used between all samples collected on each day. eDNA was collected along a transect spanning the width of the river for each replicate sample. Once the first sample was collected, the surveyor took the next sample ∼2-3m upstream, and continued this for three replicates per site. Samples were filtered on a 47-mm-diameter GF/C. Each sample consisted of filtering 1000mL of river water. Filters were placed into separate coin envelopes, which were then placed in Ziploc bags with silicone desiccant beads and stored in a freezer.

On each of the 10 days of sampling days, a negative field control was collected consisting of 500mL of distilled water being poured into a clean Nalgene in the field and processed along with field samples. Sampling was always conducted in the upstream direction, to avoid potential contamination and sediment disturbance. Field staff used new gloves between each sample, and all equipment was washed with 30% bleach following each day of field collection.

### Laboratory Processing

All filters were shipped on ice to Cramer Fish Science Genidaqs (Sacramento, California, USA, https://genidaqs.com) for DNA extraction and metabarcoding using Illumina MiSeq processing.

The DNA was extracted from each filter using a modified Qiagen DNeasy® Blood and Tissue Kit protocol. Each filter was processed overnight at 56 °C in 540 µl ATL and 60 µl Proteinase K. The resulting supernatant was passed through a Qiashredder spin column, mixed with 600 µl AL and incubated at 70 °C for 10 min. After adding 600 µl ethanol, the resulting mixture was loaded onto a DNeasy Spin column following manufacture’s protocol, with a final elution volume of 100 µl. The DNA was further processed with a Zymo Research One Step PCR Inhibitor Removal kit (Zymo Research, Irvine, California, USA). A negative control was simultaneously extracted to test for possible laboratory contamination.

Each water sample was amplified for a ∼175 base pair (bp) fragment of the mitochondrial 16S gene region which has previously been designed and validated for detection of freshwater mussels from eDNA samples (Prié et al. 2021; Marshall et al. 2022). Mussel eDNA was sequenced with MiSeq Illumina metabarcoding as previously described in Marshall et al. (2022). Library preparation followed a three step PCR described in O’Donnell et al. (2016). Each sample was amplified for a ∼175 bp fragment of the 16S gene region which has previously been tested for amplification of unionid mussels from eDNA samples (Marshall et al. 2022, Prié et al. 2021). Initial PCR amplification was completed for each sample in triplicate with 10 µl PCR reactions containing 4 µl extracted eDNA, 0.4µM primer, and Applied Biosystems™ TaqMan™ Environmental Master Mix 2.0. The amplifications started with an initial denaturation at 95°C for 5 min, followed by 35 cycles of 95°C for 15s, 5% ramp down to 55°C for 30s, and 72°C for 30s. Triplicate PCR products were diluted 1:10 then pooled prior to starting the Illumina adaptor and barcoding PCR processes.

The MiSeq library dual indexed paired-end sample preparation was adapted as described in Miya et al. (2015) from ‘16S metagenomic sequencing library preparation: preparing 16S ribosomal gene amplicon for the Illumina MiSeq system (Illumina part no. 15044223 Rev. B, San Diego, California, USA). A PCR process initiated the incorporation of Illumina adaptors and multiplexing barcodes using Prié et al. (2021) forward and reverse primers containing 33 or 34 base pairs of 5’ Illumina hanging tails to provide a priming site for a final PCR to incorporate barcodes and remaining base pairs of Illumina adaptors. The 12 µl PCR reaction contained 2 µl diluted pooled PCR product, 0.3 µM Illumina adaptor primers and 6 µl 1X Qiagen Plus Multiplex Master Mix. The PCR process denatured for 95°C for 5 min, 5 cycles of 98°C for 20s, 1% ramp down to 65°C for 15s, and 72°C for 15s., followed by 7 cycles of 98°C for 20s, 5% ramp down to 65°C for 15s, and 72°C for 15s. PCR product was diluted 1:10 prior to use in the barcode adaptor PCR process.

The final PCR incorporated paired-end dual indices (eight base pair barcodes) that allowed samples to be identified in the raw read data, and the p5/p7 adaptor sequences to allow the sample to bind onto the Illumina MiSeq flow cell. This final 12μl PCR reaction contained 1μl diluted product from the previous PCR, 0.3 µM forward and reverse indexed primer and 6ul 1X KAPA HiFi HotStart Ready Mix PCR Kit (Roche Diagnostics, Indianapolis, Indiana, USA). Conditions were 3 minutes of initial denaturation at 95°C, followed by 10 cycles at 98°C for 20 s, 5% ramp down to 72°C for 15 s, with a final 5 min 72°C extension. All PCRs were completed on Bio-Rad C1000 Touch Thermal Cyclers. Illumina adapted PCR products were pooled with equal volumes, then size selected (target ∼319bp) using 2% agarose gel electrophoresis. The final pool was sequenced with 2× 300 nt V3 Illumina MiSeq chemistry by loading 6.4 pmol library. An additional 20% PhiX DNA spike-in control was added to improve data quality of low base pair diversity samples. Additionally, a PCR no-template negative control was run for each library preparation step.

### Bioinformatic Processing and Taxonomic Identification

The data was processed following a bioinformatic pipeline previously described in Marshall et al. (2022). The forward and reverse primer sequences were removed from the demultiplexed sequences using the cutadapt (Martin et al. 2011) plugin within QIIME 2 (Bolyen et al. 2019). Next, sequence reads were filtered and trimmed using the denoising DADA2 (Callahan et al. 2016) plugin within QIIME 2. Based on the quality scores from the forward and reverse read files, a “truncLen” was set to 120 for the forward and 110 for the reverse read files. Using DADA2, error rates were estimated, sequences were merged and dereplicated, and any erroneous or chimeric sequences were removed. Unique sequences were then clustered into Molecular Operational Taxonomic Units (MOTUs) using the QIIME 2 vsearch de-novo with a 97.5% similarity threshold (Marshall et al. 2022, Marshall & Fleece 2025). MOTUs from unionid taxa were identified to the species-level using the Basic Local Alignment Search Tool (BLAST+, https://blast.ncbi.nlm.nih.gov/Blast.cgi; Camacho et al. 2009) against our custom database of both in-lab generated sequences and mt-16S sequences downloaded from NCBI GenBank. These MOTUs were further validated with comparisons against the complete NCBI nr database, to investigate alignment to mis-labeled sequences or species not historically within the sampling region. MOTUs that did not return a sequence match from the BLAST search were excluded, as they were considered not from unionid taxa.

Sequences were assigned to a species if they met a threshold of >97.5% identity and 100% query coverage. Furthermore, sequences that assigned to multiple species with the same BLAST e-value score were inspected and a final decision was made based on known distribution and presence within the sampled drainages. Additionally, if multiple sequences assigned to the same taxonomy, they were inspected and removed or collapsed into a single MOTU to obtain a final matrix of read counts per taxa. As the male mitotype is genetically distinct from the female mitotype (Curole & Kocher 2005), we separated sequences between female or male mitotypes to assess seasonal changes in the two mitotypes.

In metabarcoding analysis, the sequencing process can introduce a form of sample cross-contamination in which sequences from one sample are falsely detected within another sample, often termed ‘critical mistags’, ‘tag jumps’, or ‘index hopping’ (Esling et al. 2015, Richardson 2022). These sequencing artifacts generally represent a small fraction of the total sequences (Esling et al. 2015), yet they can lead to false detections in some circumstances. Therefore, the final processing of the MOTUs in this study consisted of estimating and removing mistags from the dataset following the framework outlined by Richardson (2022). Based on the number of reads per MOTU within the field and laboratory negative controls, the mistag rate was estimated for all dual-index combinations. Then this mistag rate was applied to all field samples to identify and remove potential cases of erroneous detection.

### Data Analysis

#### eDNA Detection Index

To assess the repeatability of eDNA detection, an eDNA Detection Index was applied to all species detections at both sites within each of the sampling events. The index levels are described as:

- **<u>Non-detection</u>** occurs when a species is not detected in any water sample replicates collected within a single site.
- **<u>Low repeatability</u>** occurs when a species is detected in **≤ 33%** of water replicates collected within a single site.
- **<u>Moderate repeatability</u>** occurs when a species is detected in **≤ 67% and > 33%** of water replicates collected within a single site.
- **<u>High repeatability</u>** occurs when a species is detected in **> 67%** of water replicates collected within a single site.

#### Male Mitotype Detection

Detections from eDNA was visualized with bubble plots to show the amount of eDNA detected for each male mitotype MOTU (i.e., measured as the number of sequence reads) on each sampling event from the two sites. For the bubble plots, the amount of eDNA detected for each MOTU on each sampling event was averaged across the water replicate samples collected at that site and on that date. The eDNA Detection Index was used within the bubble plots to illustrate detection across replicate water samples.

The amount of eDNA per sampling event was normalized by calculating the relative abundance (i.e., MOTU proportion) on each sampling date. The relative abundances were calculated using the mean amount of eDNA for each MOTU across the water replicate samples divided by the total male mitotype sequences. Barplots were used to visualize the assemblage proportions across the sampling season.

Non-metric multidimensional scaling was used to assess similarity in male mitotype MOTU detections across water sample replicates. Non-phylogenetic beta diversities were estimated using (a) a qualitative metric (calculated from presence/absence) – Jaccard dissimilarity) and (b) a quantitative metric (calculated from the assemblage proportions – Bray-Curtis Dissimilarity). These beta diversity metrics were used to assess the amount of variation in the detected mussel community across the sampling season. PerMANOVAs (vegan function adonis2) were used to test for differences in the amount of seasonal variation within Killbuck Creek replicates compared to the amount of seasonal variation within Walhonding River replicates. With homogeneity of multivariate dispersions calculated using the betadisper function and comparisons statistically tested with ANOVA. These beta diversity metrics were compared between the female and male mitotype MOTUs.

The amount eDNA detected for each male mitotype MOTU at the two sites was plotted across sampling dates. The seasonal trend in the amount of eDNA and the relative eDNA abundance for each MOTU was assessed using a regression spline in R with the *ss* function. The amount of eDNA for each MOTU on each sampling date was plotted against the gage height of the nearest USGS gage (Table 1). For male MOTUs that were able to be identified to a species-level, seasonal patterns were directly compared between male and female eDNA signal by plotting the amount of eDNA sequence read abundance across sampling events. Additionally, a male to female ratio was used to assess trends in the relationship between the male and female mitotypes across sampling timepoints. This ratio was obtained by calculating the proportion of the male mitotype compared to its female mitotype counterpart.

- **Male:female ratio**: M sequence abundance / (M sequence abundance + F sequence abundance)
- A ratio of greater than 0.5 indicates a male mitotype dominated sample, and a ratio less than 0.5 indicates a female mitotype dominated sample.

### Probability of eDNA Detection

Single-season occupancy models (MacKenzie et al. 2002) were evaluated for all male mitotype MOTUs that were identified to the species-level, to estimate the mean detection probability (*p*) (i.e., the probability of successful eDNA detection of a MOTU within a replicate environmental sample). Next, the survey design was evaluated by calculating the cumulative site-level detection probability (*p*\*) for each MOTU: *p*\* = 1-(1-*p*)^n^ where *p* is the estimated detection for a single replicate environmental sample and n is the total number of replicates collected. Occupancy models were analyzed in the R package ‘unmarked’ (Fiske & Chandler 2010). The estimated probability of eDNA detection for each MOTU was compared between detection of male and female mitotypes for mussel species that were detected with both.

## Results

The metabarcoding processing resulted in 10,022,412 raw sequence reads, of which a total of 3,496,043 reads were classified as a unionid male mitotype 16S sequence, with an average of 55,492.75 ± 54,089.11 (standard deviation) unionid male mitotype reads per sample.

Across the 10 field controls and the four laboratory controls, the number of male mussel sequences ranged from 0 to 126 total reads per sample. A total of five MOTUs occurred within the control samples, ranging from 2 to 147 total sequence reads per MOTU. The control samples consisted of 14 total occurrences of a MOTU (mean reads per occurrence = 20.9 ± 29.3 std). The mistag analysis removed all occurrences from the control samples. The mistag analysis removed a total of 26 MOTU occurrences from the 63 field eDNA samples.

### Male Mitotype Detections

A total of 24 male MOTUs were detected across the sampling events in Walhonding River and Killbuck Creek (Table 1). This included five MOTUs within Anodontini tribe, eight within Lampsilini, five within Pluerobemini, and six within Quadrulini. Of the 24 male MOTUs, only 10 were identified to the species-level, which included *Alasmodonta viridis*, *Simpsonaias ambigua*, *Lasmigona costata*, *Lasmigona complanata*, *Pygonodon grandis*, *Lampsilis siliquoidea*, *Ortmanniana ligamentina*, *Potamilus alatus*, *Fusconaia flava*, and *Quardula quadrula*. The remaining male MOTUs ranged in percent genetic identity from 86.33 to 93.89% to a species with a reference genetic sequence for the male mitotype (Table 2, Supplementary Figure 1).

**Table 2.**
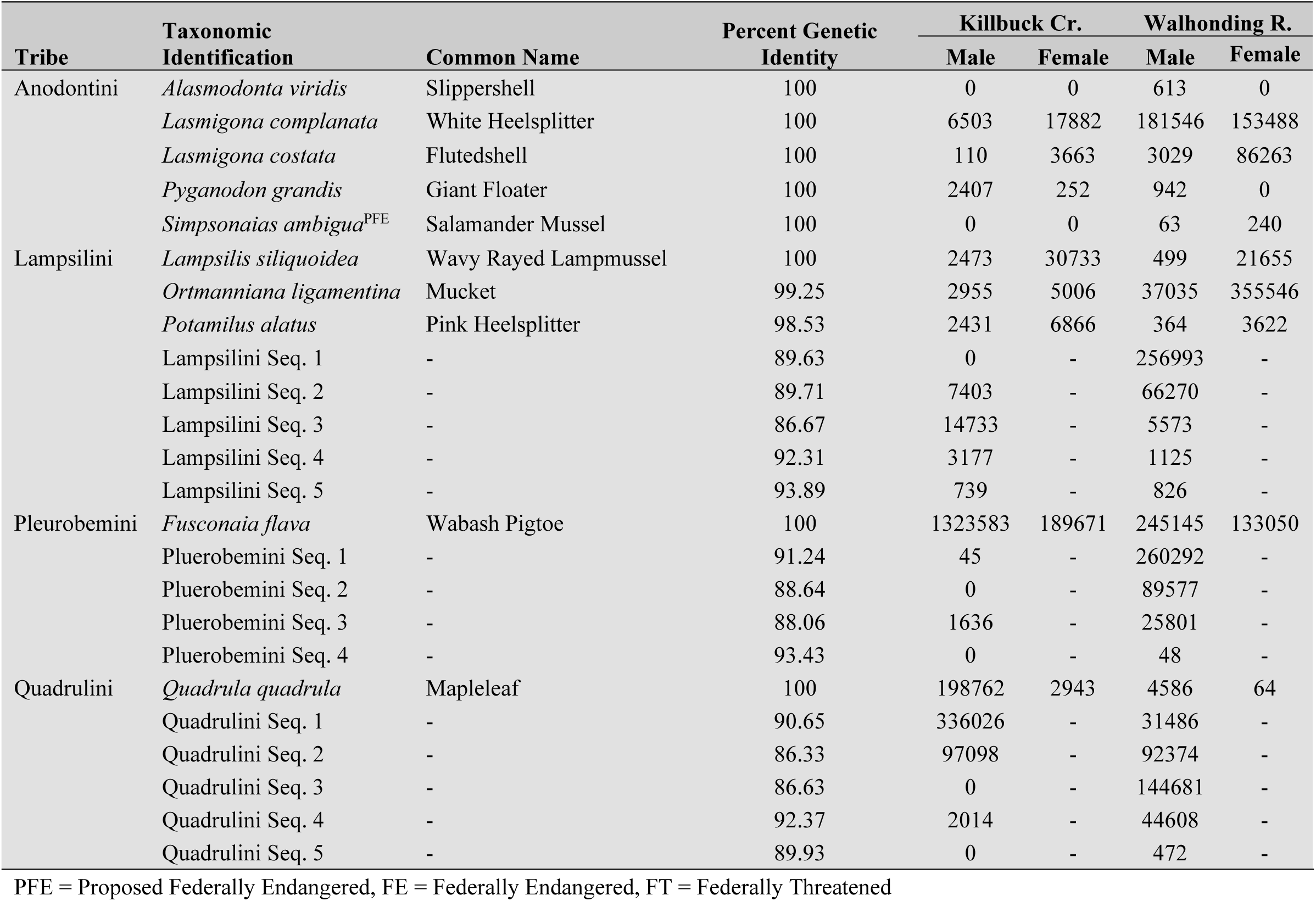
Male mitotype molecular operational taxonomic units (MOTUs) detected with environmental DNA in Killbuck Creek and Walhonding River. Comparison of environmental DNA sequences detected across the entire sampling period for the female and male mitotypes are listed for each MOTU that was discerned to the species-level.

| Tribe | Taxonomic Identification | Common Name | Percent Genetic Identity | Killbuck Cr. |  | Walhonding R. |  |
| --- | --- | --- | --- | --- | --- | --- | --- |
|  |  |  |  | Male | Female | Male | Female |
| Anodontini | *Alasmodonta viridis* | Slippershell | 100 | 0 | 0 | 613 | 0 |
|  | *Lasmigona complanata* | White Heelsplitter | 100 | 6503 | 17882 | 181546 | 153488 |
|  | *Lasmigona costata* | Flutedshell | 100 | 110 | 3663 | 3029 | 86263 |
|  | *Pyganodon grandis* | Giant Floater | 100 | 2407 | 252 | 942 | 0 |
|  | *Simpsonaias ambigua*<sup>PFE</sup> | Salamander Mussel | 100 | 0 | 0 | 63 | 240 |
| Lampsilini | *Lampsilis siliquioidea* | Wavy Rayed Lampmussel | 100 | 2473 | 30733 | 499 | 21655 |
|  | *Ortmanniana ligamentina* | Mucket | 99.25 | 2955 | 5006 | 37035 | 355546 |
|  | *Potamilus alatus* | Pink Heelsplitter | 98.53 | 2431 | 6866 | 364 | 3622 |
|  | Lampsilini Seq. 1 | - | 89.63 | 0 | - | 256993 | - |
|  | Lampsilini Seq. 2 | - | 89.71 | 7403 | - | 66270 | - |
|  | Lampsilini Seq. 3 | - | 86.67 | 14733 | - | 5573 | - |
|  | Lampsilini Seq. 4 | - | 92.31 | 3177 | - | 1125 | - |
|  | Lampsilini Seq. 5 | - | 93.89 | 739 | - | 826 | - |
| Pleurobemini | *Fusconaia flava* | Wabash Pigtoe | 100 | 1323583 | 189671 | 245145 | 133050 |
|  | Pluerobemini Seq. 1 | - | 91.24 | 45 | - | 260292 | - |
|  | Pluerobemini Seq. 2 | - | 88.64 | 0 | - | 89577 | - |
|  | Pluerobemini Seq. 3 | - | 88.06 | 1636 | - | 25801 | - |
|  | Pluerobemini Seq. 4 | - | 93.43 | 0 | - | 48 | - |
| Quadrulini | *Quadrula quadrula* | Mapleleaf | 100 | 198762 | 2943 | 4586 | 64 |
|  | Quadrulini Seq. 1 | - | 90.65 | 336026 | - | 31486 | - |
|  | Quadrulini Seq. 2 | - | 86.33 | 97098 | - | 92374 | - |
|  | Quadrulini Seq. 3 | - | 86.63 | 0 | - | 144681 | - |
|  | Quadrulini Seq. 4 | - | 92.37 | 2014 | - | 44608 | - |
|  | Quadrulini Seq. 5 | - | 89.93 | 0 | - | 472 | - |

PFE = Proposed Federally Endangered, FE = Federally Endangered, FT = Federally Threatened

Within Killbuck Creek, 17 total male MOTUs were detected across the sampling season, which ranged from 45 sequence reads (Pluerobemini Seq. 1) to 1,323,583 (*Fusconaia flava*) (Table 2). All of the 24 male MOTUs were detected in Walhonding River, which ranged from 48 (Pluerobemini S. 4) to 260,292 (Pluerobemini Seq. 1) (Table 2). Overall, Killbuck Creek had a higher total sequence read count for male MOTUs across the sampling season (Figure 2). Killbuck Creek was largely dominated by *Fusconaia flava*, whereas Walhonding River had four male MOTUs that ranged from 12 to 17% of the total sequence read counts detected (Table 2, Figure 2).

**Figure 2.**
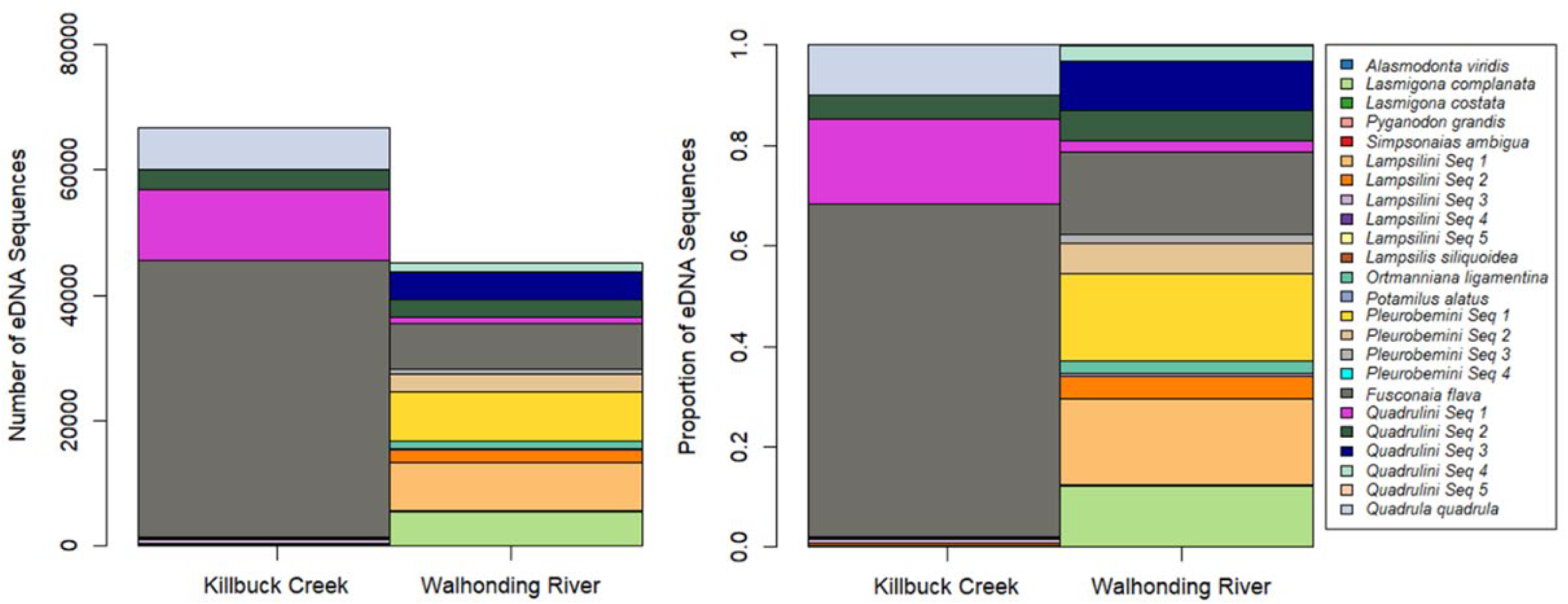
(A) Mean number of freshwater mussel environmental DNA sequences per sample and (B) proportion of freshwater mussel environmental DNA sequences detected in Killbuck Creek and Walhonding River

### Seasonal Trends

The majority of detections in both Killbuck Creek and Walhonding River were classified as high repeatability (Figure 3). There were significant differences between the amount of DNA sequence reads obtained at the three levels of the eDNA detection index, where detections occurring at high repeatability had on average the greatest amount of sequence reads, and detections at moderate and low repeatability had on average similar amounts of sequence reads (ANOVA: F-value = 99.91, *p* < 0.001) (Figure 3).

**Figure 3.**
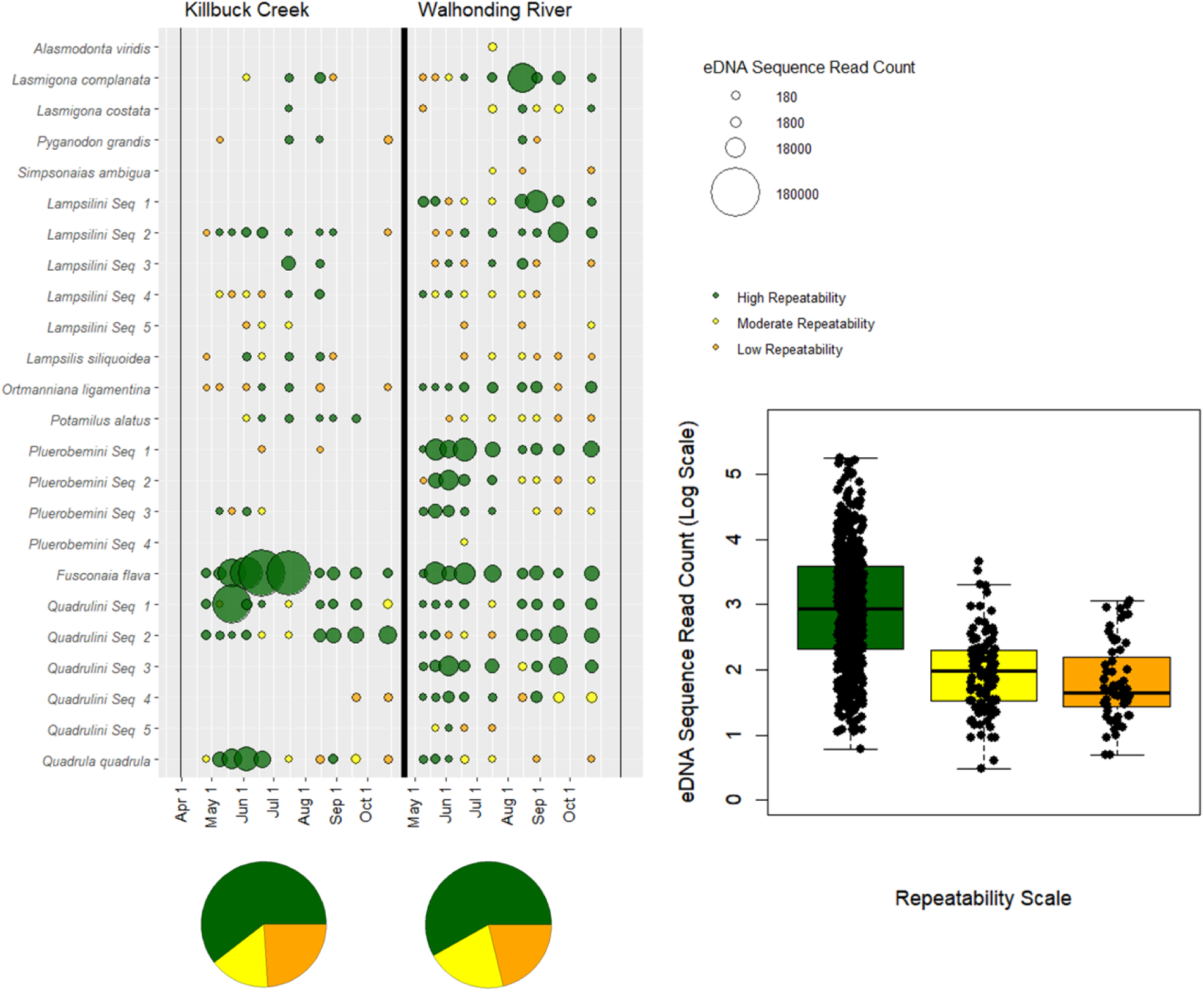
Bubble plot of seasonal environmental DNA detections for freshwater mussels in Killbuck Creek and Walhonding River. Size of bubbles represent the mean eDNA sequence and colors represent repeatability of detection across replicates

The relative eDNA abundance estimates of MOTU male mitotypes varied across sampling season in both Killbuck Creek and Walhonding River (Figure 4). Notably, within Killbuck Creek, the male mitotype sequence read abundance was far greatest from May thru July compared to early spring or later summer sampling events (Figure 4). Seasonal trends in sequence read abundance can be found in Supplementary Figure 2 & Supplementary Figure 3. Community assemblage displayed seasonal fluctuations in both Killbuck Creek and Walhonding River (Figure 4, Supplementary Figure 4).

**Figure 4.**
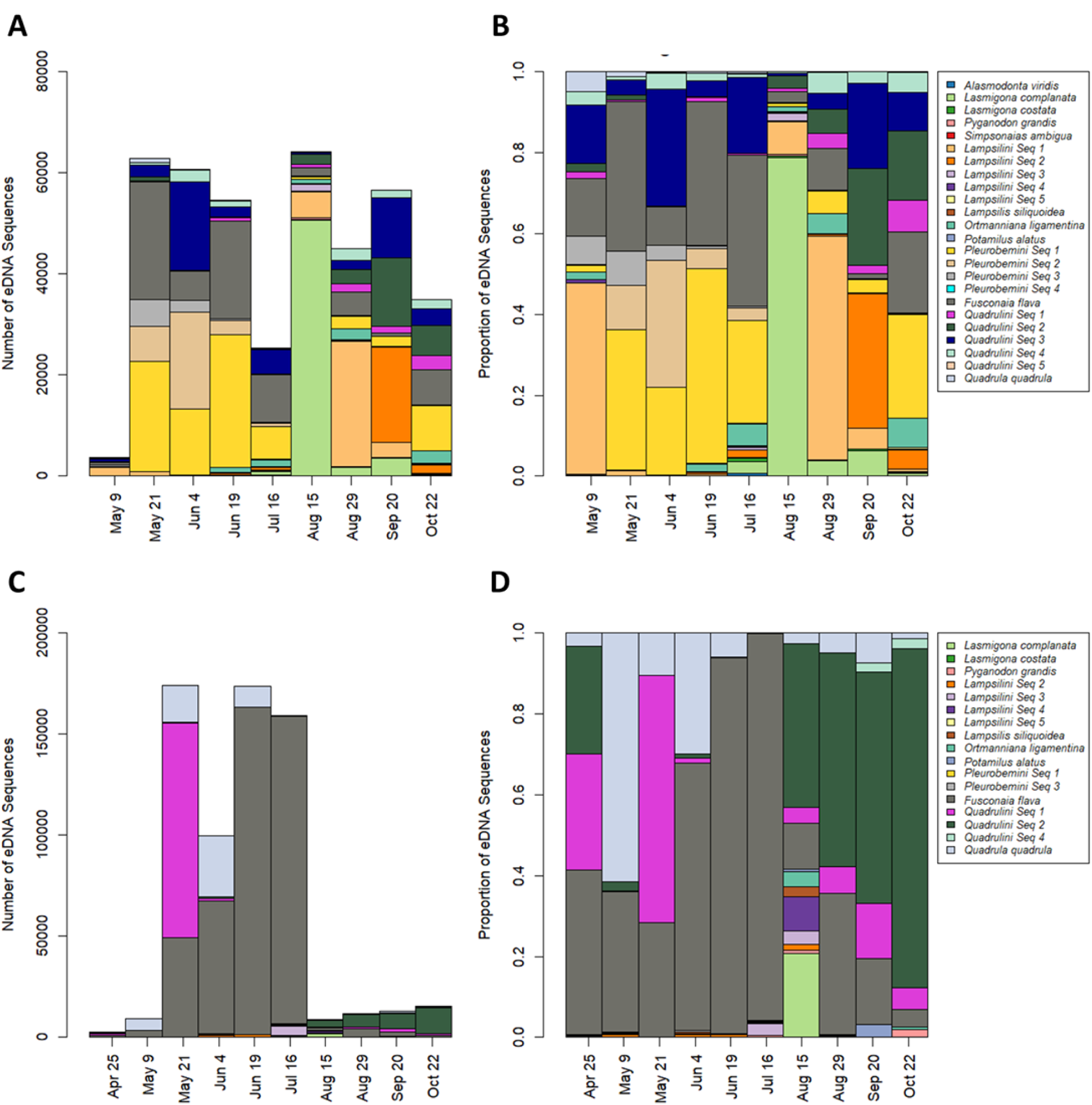
(A) Number of freshwater mussel environmental DNA sequences and (B) proportion of freshwater mussel environmental DNA sequences detected across sampling events in Walhonding River. (C) Number of freshwater mussel environmental DNA sequences and (D) proportion of freshwater mussel environmental DNA sequences detected across sampling events in Killbuck Creek.

Beta diversity estimates (i.e., Bray Curtis Dissimilarity and Jaccard Dissimilarity) and NMDS analysis were used to further assess patterns in the male mitotype assemblage over the sampling period. Patterns within both beta diversity metrics were similar, where the samples clustered within their respective sampling site (i.e., Killbuck Creek vs Walhonding River) (Supplementary Figure 4, circles vs squares). Additionally, samples showed low variation across sampling replicates collected on the same sampling date (Supplementary Figure 4 & Supplementary Figure 5).

To further assess seasonal fluctuations in the detected male mitotype assemblage, the beta diversity metrics were assessed against the amount of time between the sampling events and across flow exceedances. In both Killbuck Creek and Walhonding River, the community dissimilarity increased with increasing number of days between sampling events (Supplementary Figure 6). Notably, the mean variance in community dissimilarity across the sampling season was greater for the male mitotype than the female mitotype within both Killbuck Creek and Walhonding River (see boxplots in Supplementary Figure 6)

### Comparisons of Female and Male Mitotypes

Of the nine male MOTUs that were able to be identified to a species, all but one were also detected in eDNA with the female mitotype. *Alasmodonta viridis* is the lone species detected solely with the male mitotype where it was detected with only 613 sequence reads in the Walhonding River (0.04% of the total male mitotype eDNA in the Walhonding River) (Table 2). While *Pyganodon grandis* was detected at both sites within the male mitotype eDNA it was only detected with female mitotype eDNA from the Killbuck Creek (Table 2). Conversely, one species detected with female mitotype eDNA that does have a male 16S reference sequence but was not detected within the male mitotype eDNA - *Ptychobranchus fasciolaris*. This species only accounted for less than 0.03% of the total female mitotype eDNA in either Walhonding River or Killbuck Creek (464 total female eDNA sequence reads for *P. fasciolaris*, and thus represented a very rare detection. The rest of the male MOTUs represent species with missing male 16S reference sequences, and thus it is unknown exactly which species from the female mitotype eDNA dataset they correspond to.

Species identified with the male mitotype were further assessed by comparing their male mitotype detection trends with documents of known gravidity recorded in Watters et al. (2009). Spawning is assumed to occur in the period prior to the of gravid observation females, and indeed the current study found that peaks in male mitotype eDNA typically coincided with the period leading to known gravidity (Figure 5). For example, *Alasmidonta viridis* is expected to spawn in June or July as it is known to be gravid from August through May. Subsequently, this study found the only trace of male mitotype eDNA form *A. viridis* occurred in July. Similar trends were observed for *S. ambigua, L. complanata, L. costata, P. grandis, O. ligamentina,* and *F. flava* (Figure 5). Furthermore, the ratio of male to female eDNA provided a direct assessment of spikes in male mitotype signal (Figure 5).

**Figure 5.**
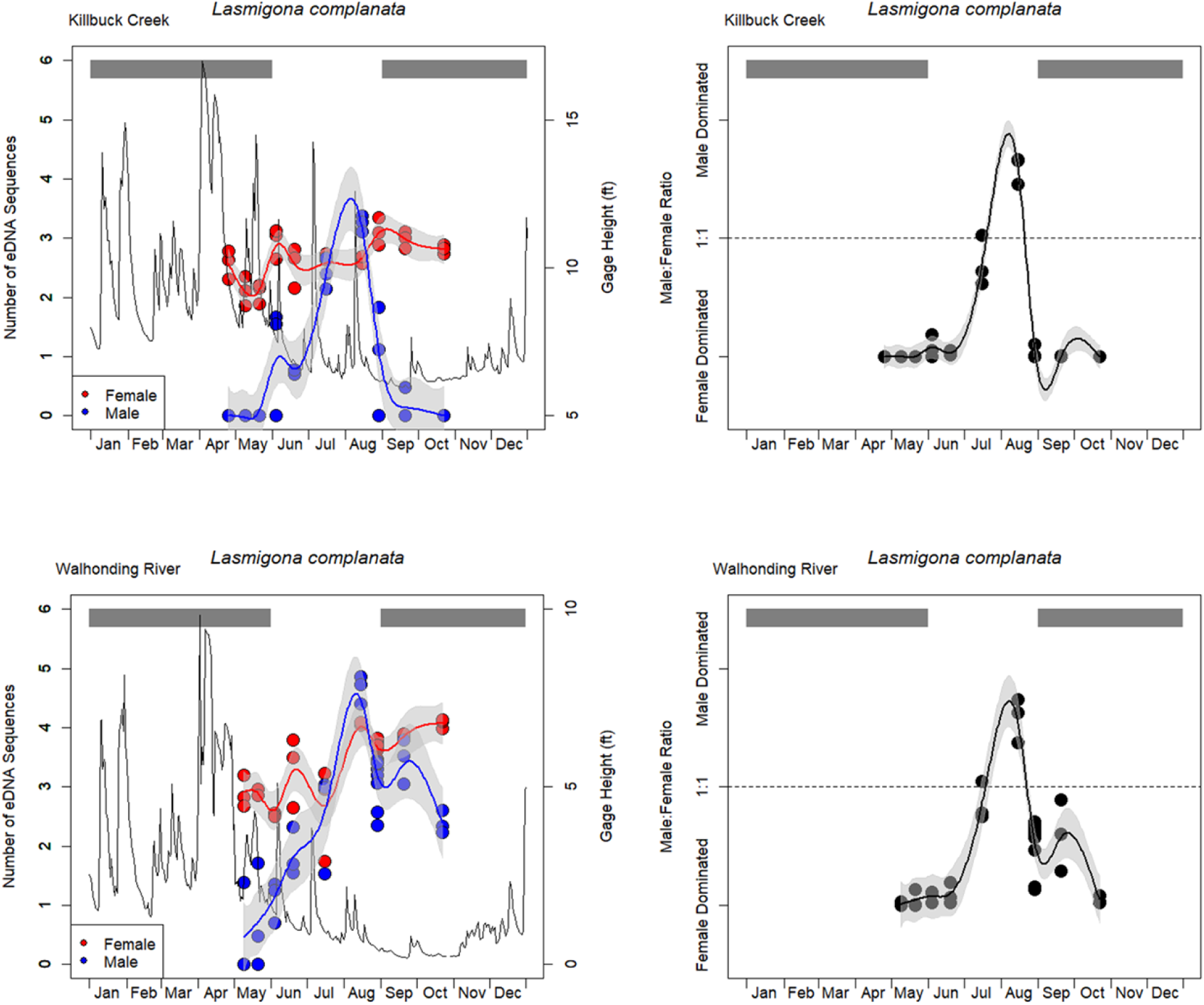
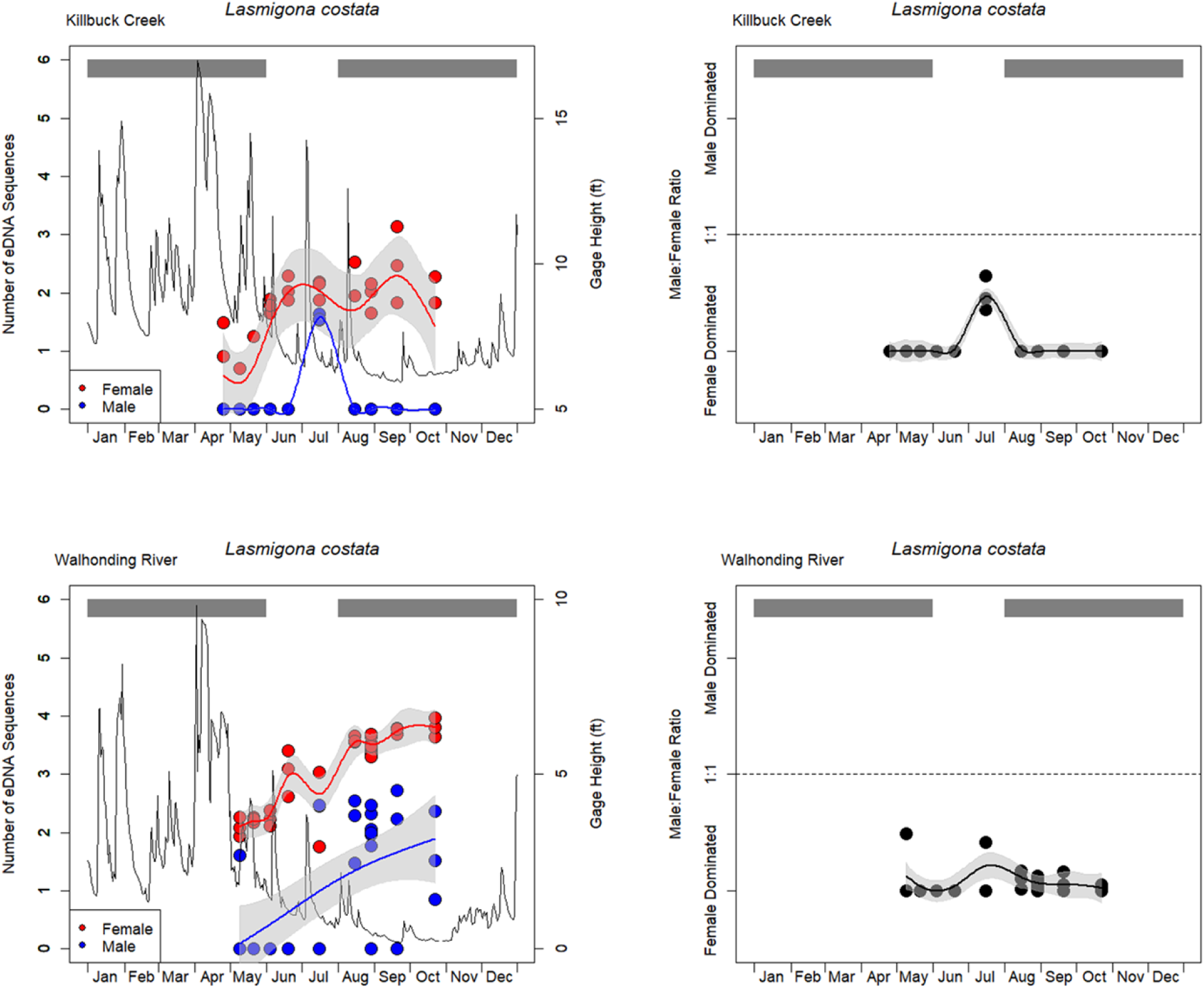
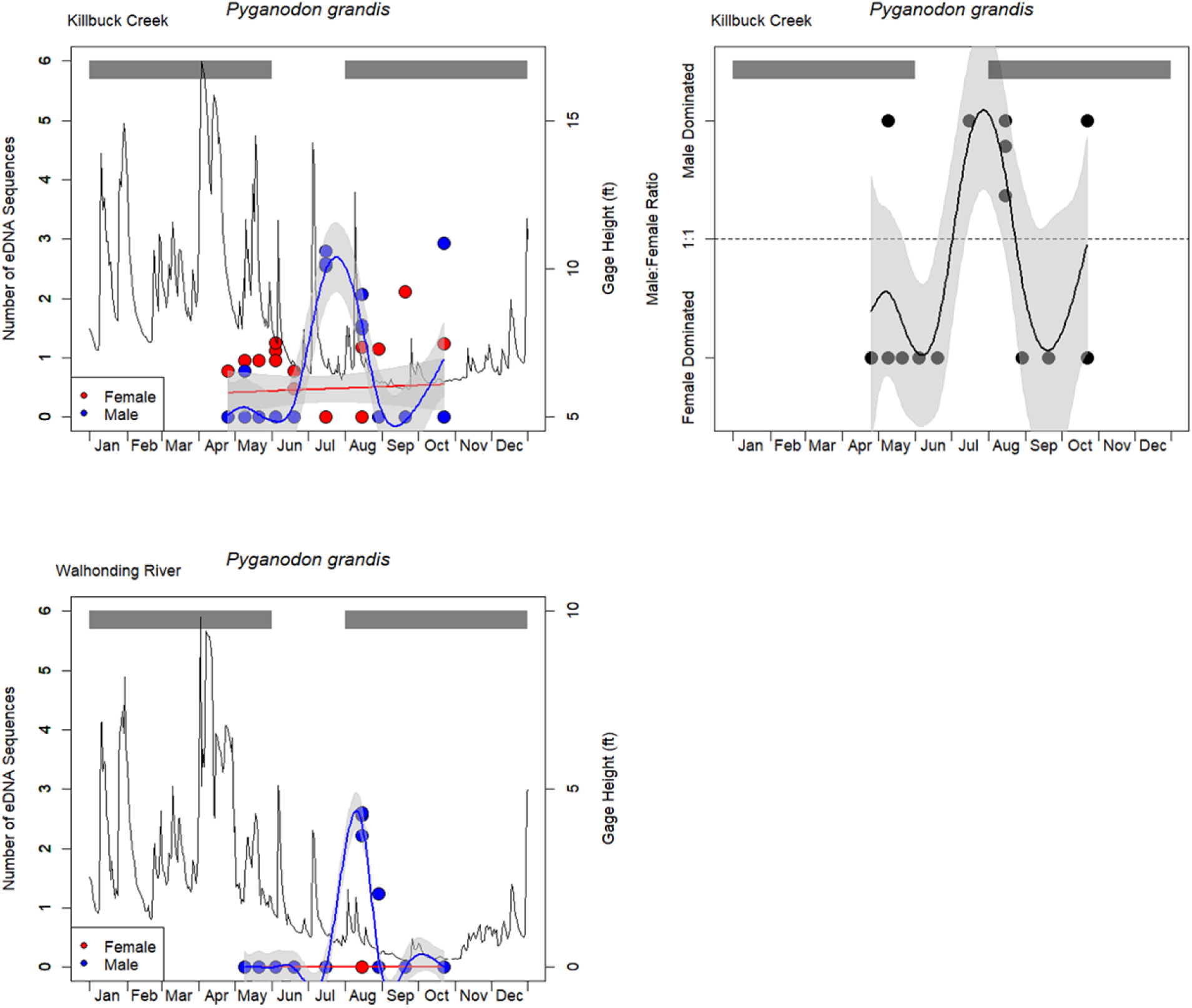
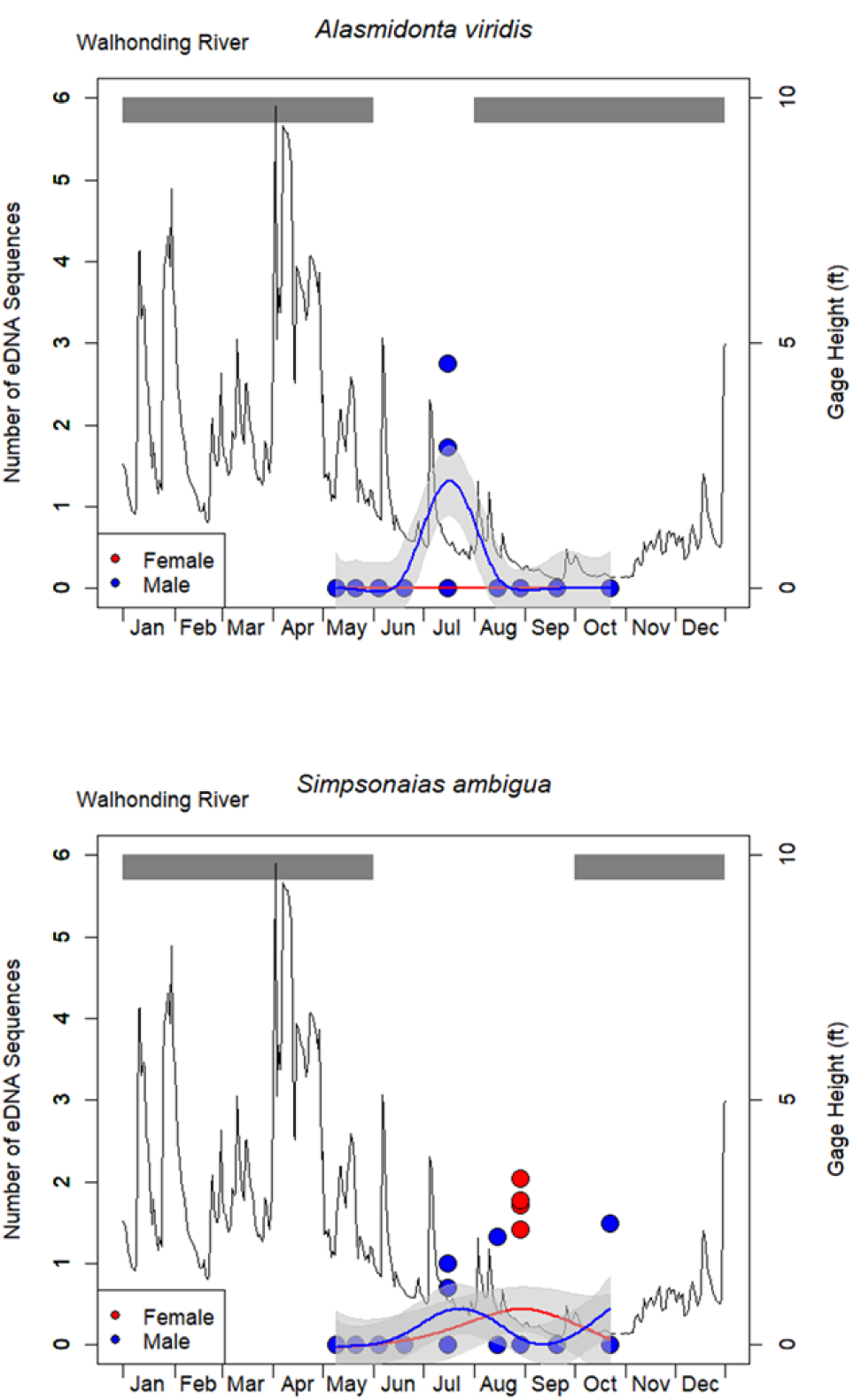
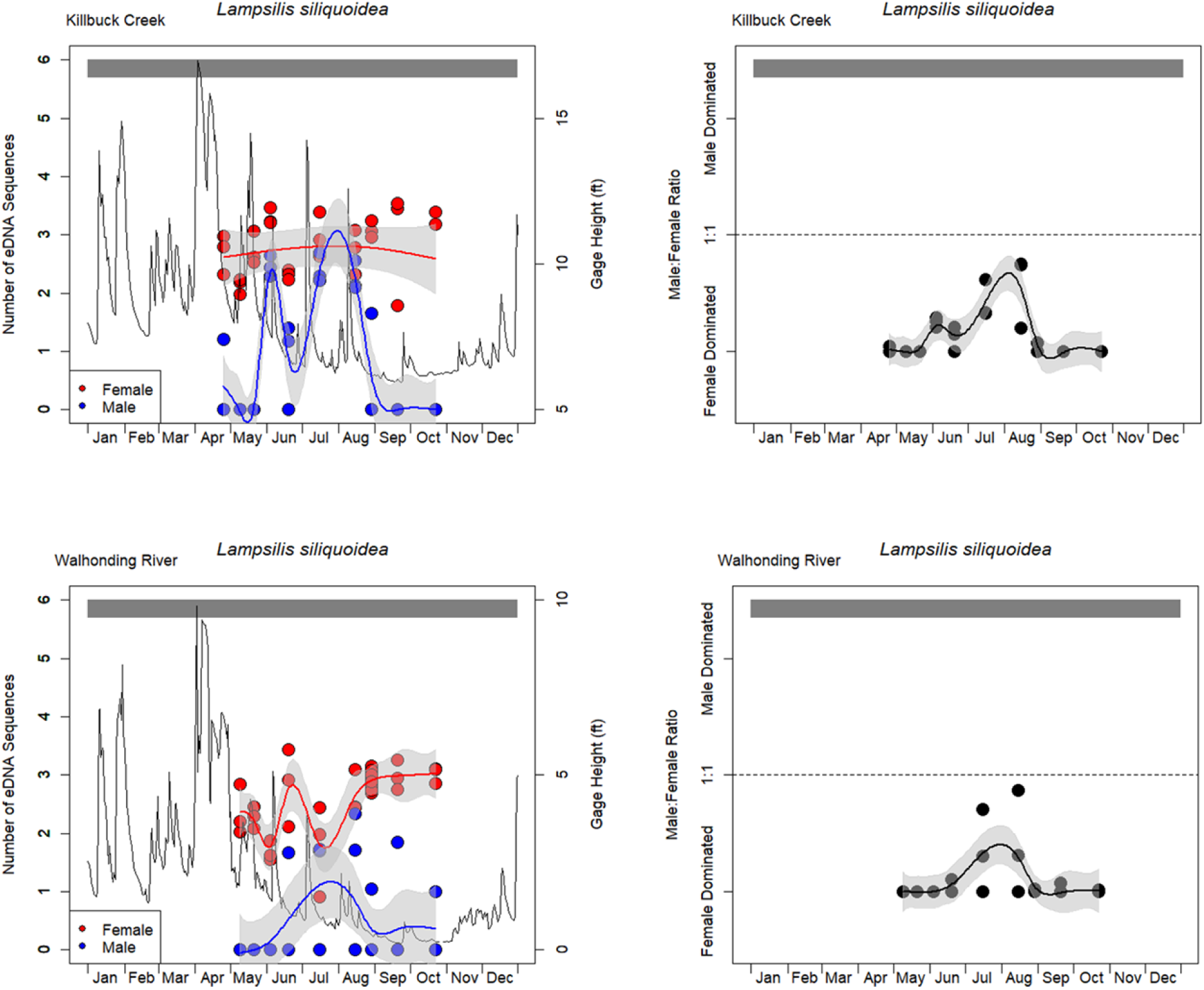
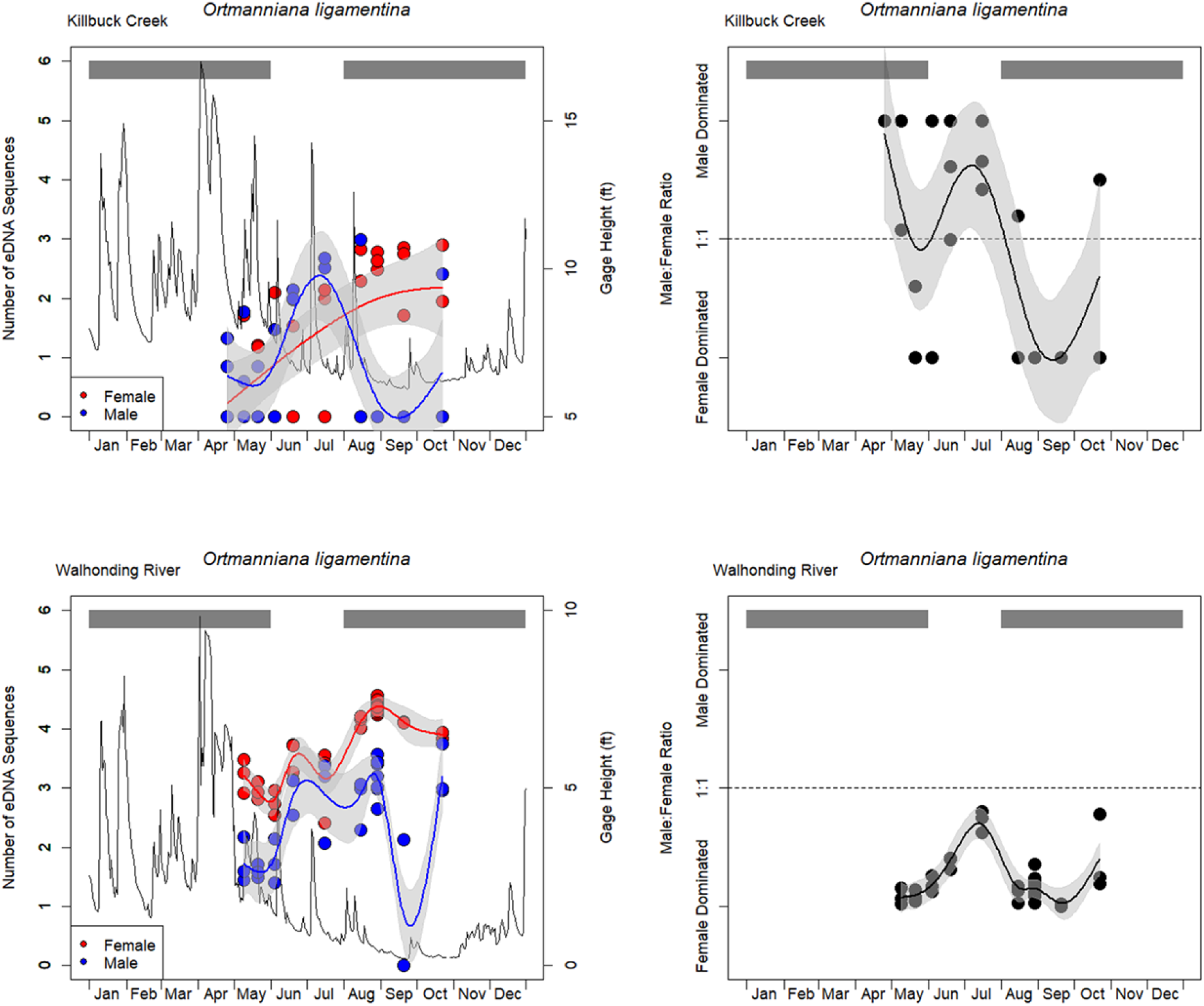
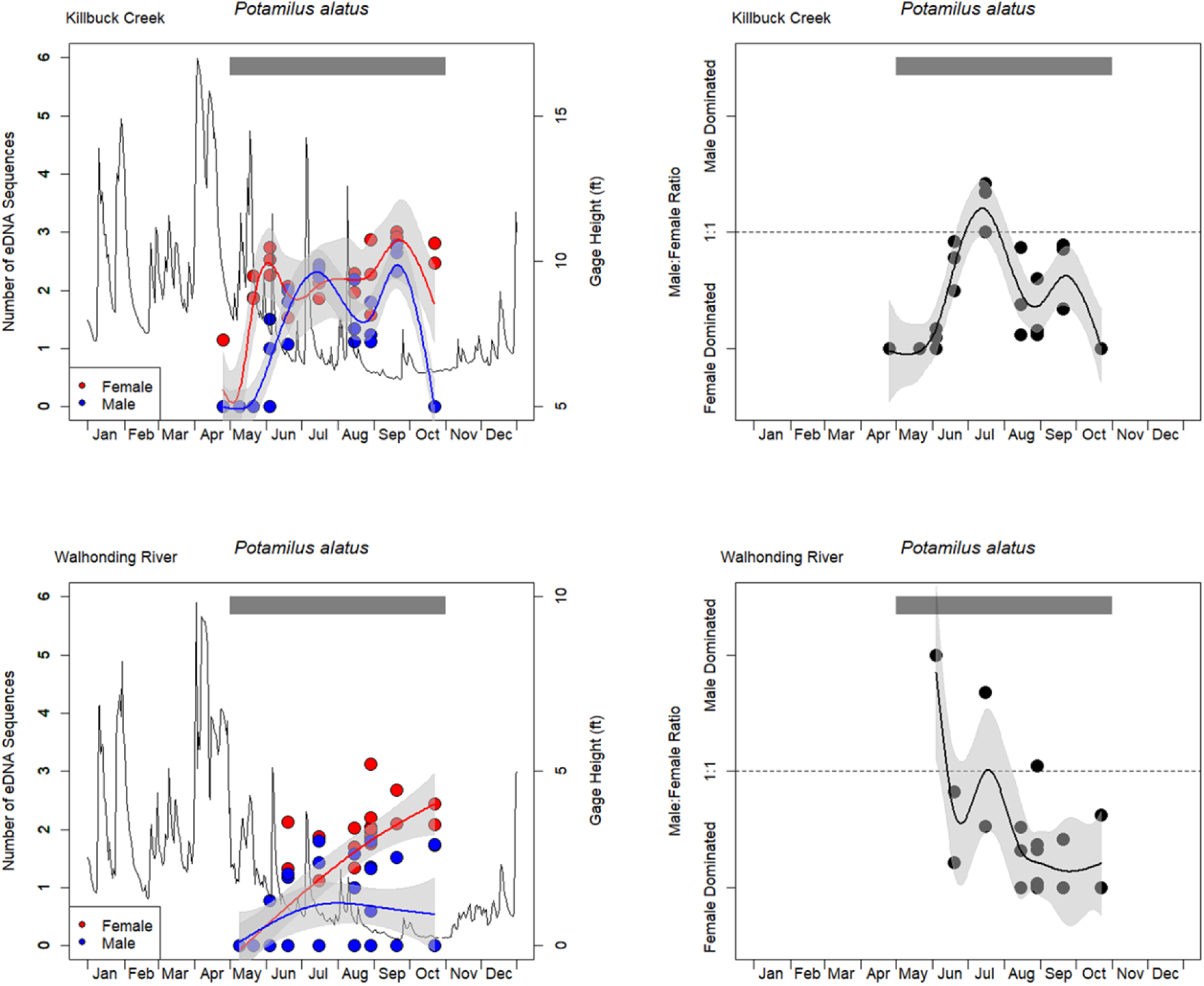
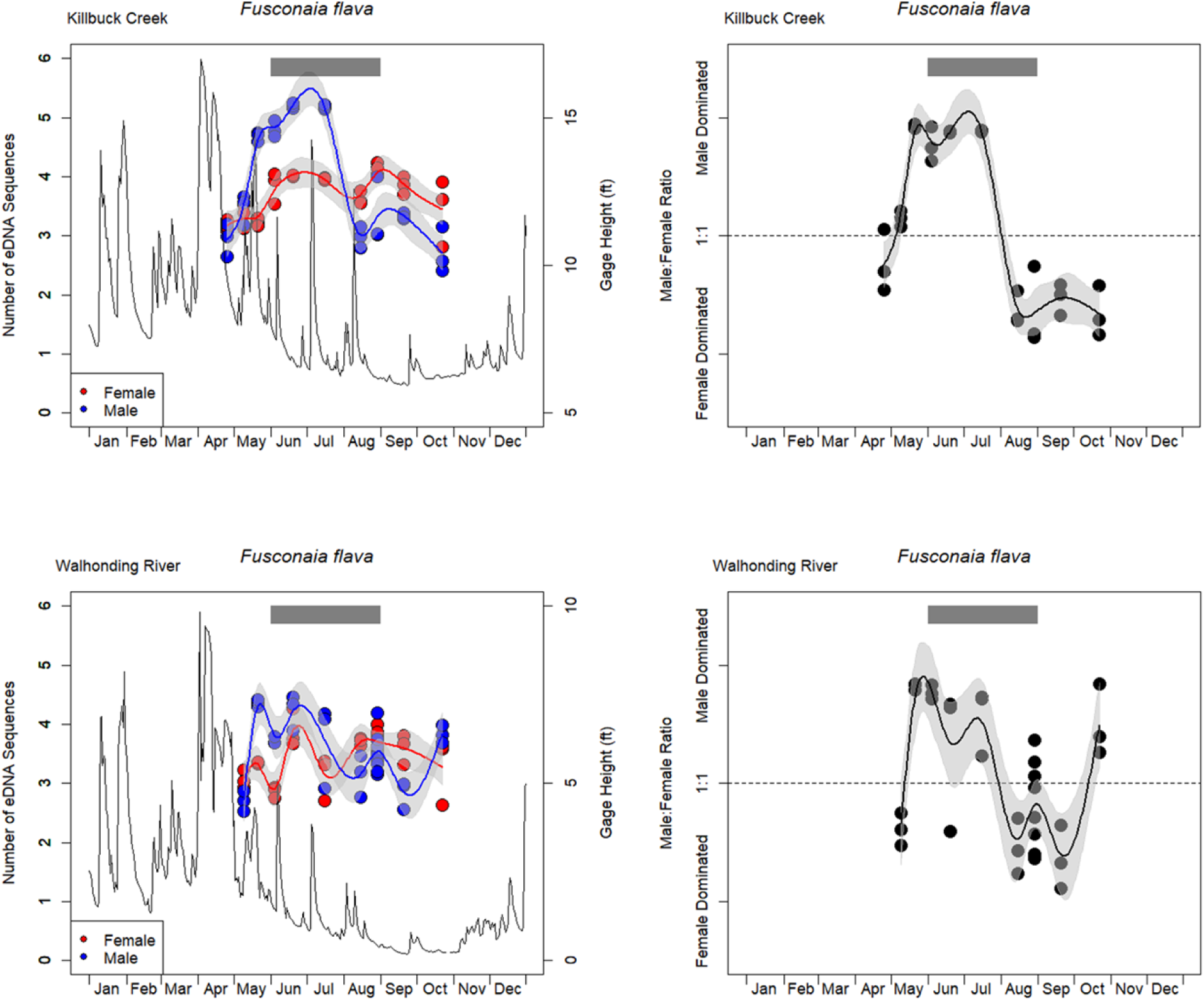
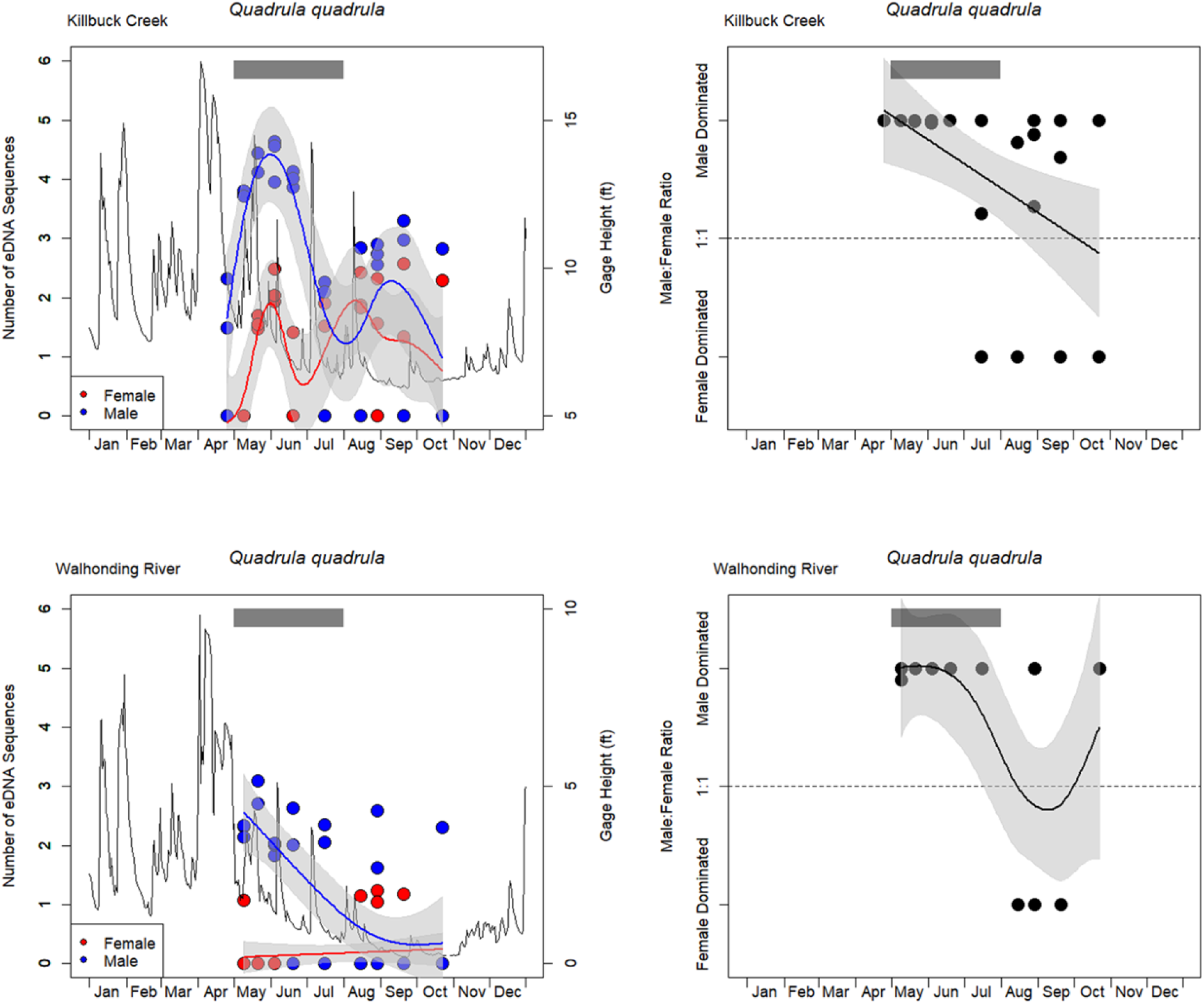
Trends in the environmental DNA sequence abundance for the female and male mitotypes and the male to female ratio across sampling points in Killbuck Creek and Walhonding River. Plots are shown for each species that was discerned with the male mitotype at either sampling site. Grey horizontal bars at the top of each plot illustrate periods of gravidity in the literature (Watters et al. 2009). Black trend lines represent river height from nearest USGS gage.

Nine species detected with both the male and female mitotypes were directly compared across their eDNA detection probability. Six of the eight species displayed a higher detection probability for the female mitotype compared to their male mitotype (Figure 6). *Fusconaia flava* was detected in all samples with both mitotypes and thus had a 0.99 probability of eDNA detection with either mitotype (Figure 6). *Quadrula quadrula* was the lone species to have a higher estimated detection probability with the male mitotype compared to its female counterpart (Figure 6).

**Figure 6.**
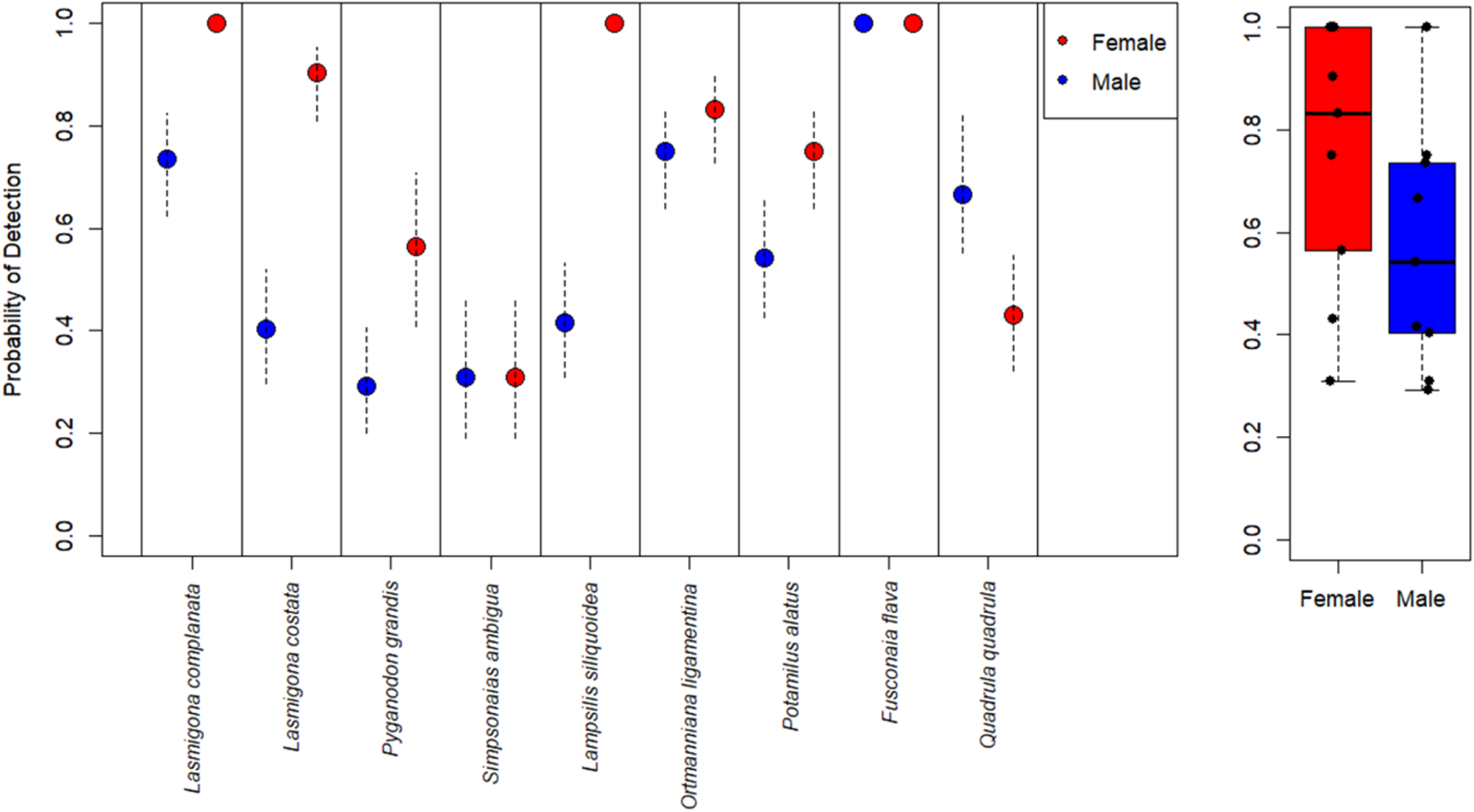
The median probability of detection estimates from environmental DNA for the female or male mitotype for each species detected with both mitotypes. Error bars represent credibility intervals for probability of detection estimates.

## Discussion

This study detected 24 MOTUs identified as freshwater mussel male mitotype from eDNA collected across seasonal sampling at two diverse mussel beds. The seasonal sampling showcased trends in male mitotype eDNA detection for several freshwater mussel species, with peaks in male mitotype eDNA typically aligning with expected spawning times based on literature records of gravidity in females (Watters et al. 2009). For example, bradytictic species such as *P. grandis*, *L. costata*, and *O. ligamentina* all had elevated signals of male mitotype in July, coinciding with the expected spawning time of long-term brooders with female gravidity occurring from early autumn until early spring (Watters et al. 2009, Haag 2012). Furthermore, many sampling events had no detection of the male mitotype for these species despite their female mitotype counterpart being detected across all sampling events, further suggesting time periods of elevated male mitotype signal are related to a reproductive behavior.

For species with direct comparisons of eDNA detection of the male mitotype to its female counterpart, the female mitotype was typically detected in a higher number of samples and subsequently the female mitotypes displayed higher probability of detection across the sampling season. This may suggest the female mitotype is more likely to be consistently shed and detected between April thru October, whereas the male mitotype may be more episodic in detection and/or linked to a seasonal behavior response. Additionally, *A. viridis* and *P. grandis* were exclusively detected with only the male mitotype in the Walhonding River. Notably, the timing of *P. grandis* male mitotype detection in the Walhonding River coincided with elevated male signals for this species in Killbuck Creek, further supporting these male mitotype detections are linked to spawning behavior.

The detection of these species exclusively through the male mitotype suggests that sperm-associated eDNA may disperse across greater distances than non-spawning eDNA sources, such as DNA derived from sloughed stomach cells during filter feeding or from metabolic waste. Ishibashi et al. (2000) reported that spermatozeugmata could remain viable for 48 to 72 hours and Fergeson et. al. (2013) reported that females could be fertilized by males as far as 15 kilometers away. Although male mitotype signals are generally less likely to be detected outside of spawning events, their presence can broaden spatial coverage and enhance assessments of regional mussel diversity, as spermatozeugmata are capable of traveling farther than other eDNA sources (Marshall et al. 2025). For instance, *S. ambigua* was previously unrecorded this far upstream in the Walhonding River (Watters et al. 2009) and was not observed during visual surveys (EDGE 2025). Nevertheless, *S. ambigua* was repeatedly detected in eDNA samples via the male mitotype from July through October, indicating its potential occurrence upstream of the current sampling location. Environmental DNA sampling has proven useful in detecting new populations of rare mussel species (Douglas et al. 2025), and detection of the male mitotype may be used as a coarse filter for characterizing whether rare and elusive mussel species remain extant at reach and basin scales.

While peaks in the eDNA signal were witnessed for most male mitotype MOTUs, several of these MOTUs were detected across the entire sampling period, and thus the mere presence of a male mitotype within an eDNA sample does not appear to be indicative of spawning on its own. Several non-sperm related sources may influence the concentration of the male mitotype within eDNA samples, which may lead to an inaccurate interpretation of a spawning event. For example, a recently deceased male individual could lead to a heightened male mitotype signal as the gonadal tissue decays. Additionally, genetic studies have found that several species have male mitotype “leakage” where it is present in low quantities within somatic tissue (Mioduchowska et al. 2016, Brenton et al. 2017). This “leakage” phenomenon could result in the detection of the male mitotype even from eDNA sources originating non-gonad sources (e.g., shed somatic tissue).

Additionally, gravid females release glochidia larvae into the environment via conglutinates, broadcast release, and lures (Haag 2012), and at certain times of the year these microscopic glochidia can be collected within the water column (Culp et al. 2011). The collection of male glochidia within an eDNA sample may also result in a heightened signal of male mitotype, despite the lack of sperm within environment. Finally, some non-unionid bivalve broadcast spawners perform post-spawning gamete clearance whereby unused or undeveloped sperm is expelled at the end of a reproductive season (Deguchi & Osada 2020). This clearance helps reset the reproductive system for the next cycle and prevents buildup of aging or non-viable gametes. It is possible freshwater mussels perform post-spawning gamete clearance, which would likely result in an elevated signal of the male mitotype.

Due to these potential sources of non-sperm related male mitotype detection, a better proxy for a spawning event may be to track the relative concentration of the male mitotype in comparison to its female counterpart. In the presence of spermatozeugmata, the concentration of the male mitotype is expected to increase relative to the female mitotype because (1) spermatozoa only consist of the male mitotype (Zouros 2013), (2) each individual spermatozoa contain several mt-genomes (Waller and Lasee 1997), and (3) each spermatozeugmata consists of thousands of spermatozoa (Waller and Lasee 1997). Therefore, applying a male-to-female ratio offers a standardized approach for identifying periods when the male mitotype occurs at higher concentrations than its female counterpart, potentially indicating the collection of spermatozeugmata within an eDNA sample.

Some species had a continuous signal of male mitotype throughout the entire sampling period, and even displayed similar seasonal eDNA trends between the female and male mitotypes (e.g., *F. flava* and *Q. quadrula*). For example, *F. flava* male mitotype was detected in every eDNA sample at both Walhonding River and Killbuck Creek, and thus it may be difficult to relate eDNA detection to spawning behavior for this species. However, the male mitotype for *F. flava* did display the greatest elevated signal from May through July, where the sequence read abundance of the male mitotype was greater than 10x that of its female counterpart at both sites during this time. While it’s unknown why there is a consistent eDNA signal of *F. flava* male mitotype throughout the year, the male dominated samples in May through July may suggest the presence of sperm in those months.

Conversely *Q. quadrula* displayed a greater signal of the male mitotype compared to the female mitotype across most timepoints from either Killbuck Creek or Walhonding River. Furthermore, *Q. quadrula* was the lone species to have a higher estimated detection probability for the male mitotype over its female counterpart. In addition to *Q. quadrula*, most male MOTUs from the Quadrulini tribe were found to be detected throughout the entire extent of the current study, and often at high sequence read abundance. This may suggest there is little to no seasonal influence in the shedding and detectability of the male mitotype for Quadrulini mussels. The Quadrulini tribe is broadly considered tachytictic with spawning occurring in early to late spring (Watters et al. 2009, Sietman et al. 2012, Haag 2012). However, *Quadrula fragosa* (Winged Mapleleaf) was recently noted as an outlier within the Quadrulini, with spawning occurring in late summer (Hove et al. 2012, Sietman et al. 2012). Regardless of the exact spawning period, no Quadrulini has ever been documented as exhibiting multiple spawning events throughout a year (Watters et al. 2009, Sietman et al. 2012). Therefore, it is unclear why the male mitotype appears to be highly concentrated and detectable from May through October for most of the Quadrulini mussels in the current study. Further work is required to investigate how male mitotype eDNA release may differ across mussel tribes and species, which may complicate interpretation of spawning events.

Complex population demographics and evolutionary life history traits, such as hermaphroditism, are likely to influence the presence and detection of the male mitotype within eDNA. Complete hermaphroditism has been identified in a few North American freshwater mussels, which includes two species present within the current study – *Utterbackia imbecillis* and *Lasmigona compressa* (van der Schalie 1970). These complete hermaphroditic species have entirely lost the unique male mitotype found within freshwater male gonad tissue, and instead all tissue now possess only the female mitotype (Guerra et al. 2019). Therefore, the presence of sperm will not be detectable through the analysis of male mitotype eDNA for these few species.

While only a few species have been identified as complete hermaphrodites, several others have been identified as sporadic hermaphrodites where a small proportion of the population is monoecious (van der Schalie 1970). This includes five species present at the sampling sites in the current study – *Fusconaia flava, Quadrula quadrula, Eyrunia dillatata, Ptychobranchus fasciolaris*, *Pyganodon grandis*, and *Lasmigona complanata*. While some individuals are hermaphroditic, the population still retains both the female and male mitotypes. Of the five species previously identified as having sporadic hermaphrodism, *F. flava* and *Q. quadrula* both had relatively strong signal of male mitotype on all sampling dates from May through October. Future assessments of spawning behavior will benefit from understanding population demographics, such as frequency of sporadic hermaphrodism, to better understand how this may influence shedding and detection rates of male mitotype eDNA.

The male mitotype is not often sequenced for phylogenetic or eDNA based studies, and thus most MOTUs for the male mitotype cannot be discerned to a species based on current reference genetic repositories. A more complete database would go a long way to improving our understanding how to interpret male mitotype eDNA patterns. Yet, this study demonstrates that male mitotype eDNA can provide valuable insights into freshwater mussel reproductive ecology, though interpretation requires caution. Peaks in male mitotype detection often aligned with expected spawning periods, supporting its potential as a proxy for sperm release. However, continuous or sporadic signals of male mitotype may highlight the influence of non-spawning sources, population demographics, and life history traits such as hermaphroditism. The consistent detection of female mitotypes underscores their reliability as a baseline, while male-to-female ratios may offer a more refined approach for describing spawning events. These findings establish male mitotype eDNA as a promising, non-invasive tool which could be a useful technique for (1) determining when to look for gravid females, (2) providing new insight into understudied reproductive behaviors, and 3) investigating environmental cues that trigger reproductive behaviors (e.g., temperature).

## Acknowledgments

This project was supported through field support during eDNA surveys by Charlie Allen, Zoe True and Ashley Hansen from Stantec.

## Data Availability Statement

Supplementary data files are provided to Open Science Framework at https://doi.org/XXXX. Raw sequence data are deposited in the National Center for Biotechnology Information (NCBI) Sequence Read Archive (SRA) under the BioProject accession XXXXX.

## Ethics and Permit Approval Statement

No permitting was required for eDNA surveying for this project.

## Conflict of Interest Disclosure

The authors declare no conflicts of interest.

Permission to Reproduce Material from other sources: All material is original to this manuscript.

---

**Supplementary Figure 1.**
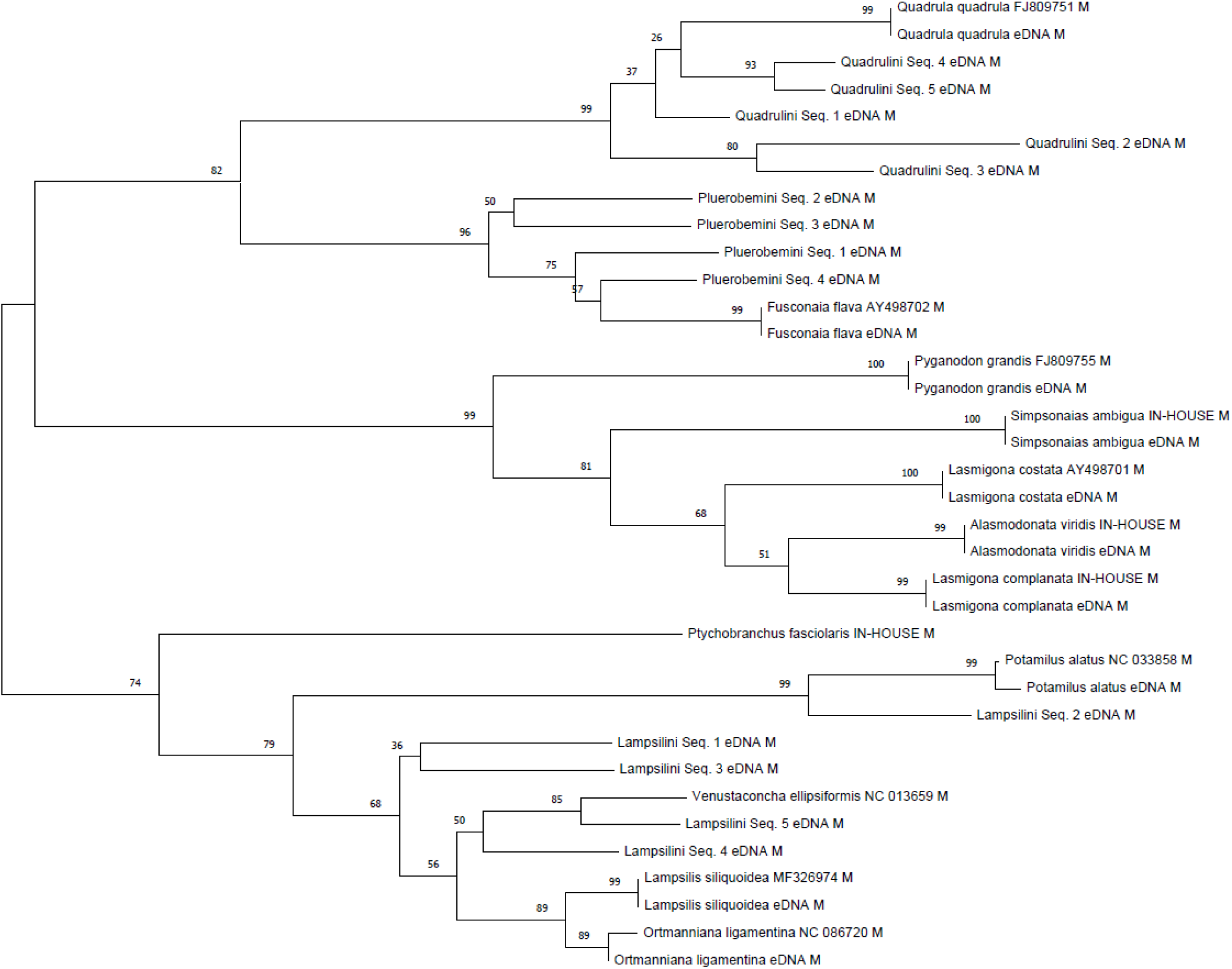
Phylogenetic tree used for taxonomic identification of the mussel male mitotype environmental DNA sequences based on the genetic match to known genetic data within the NCBI Genbank repository

**Supplementary Figure 2.**
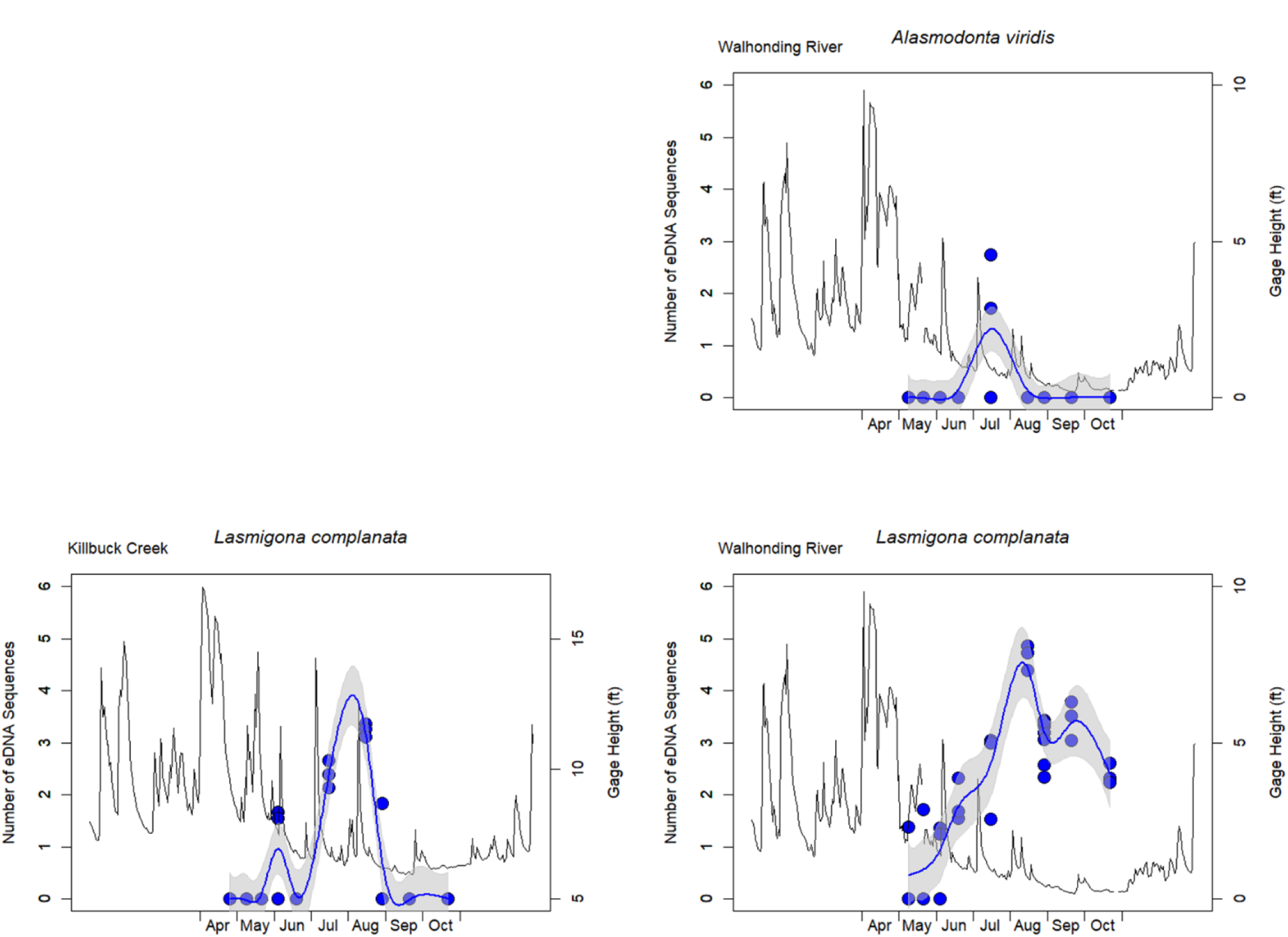
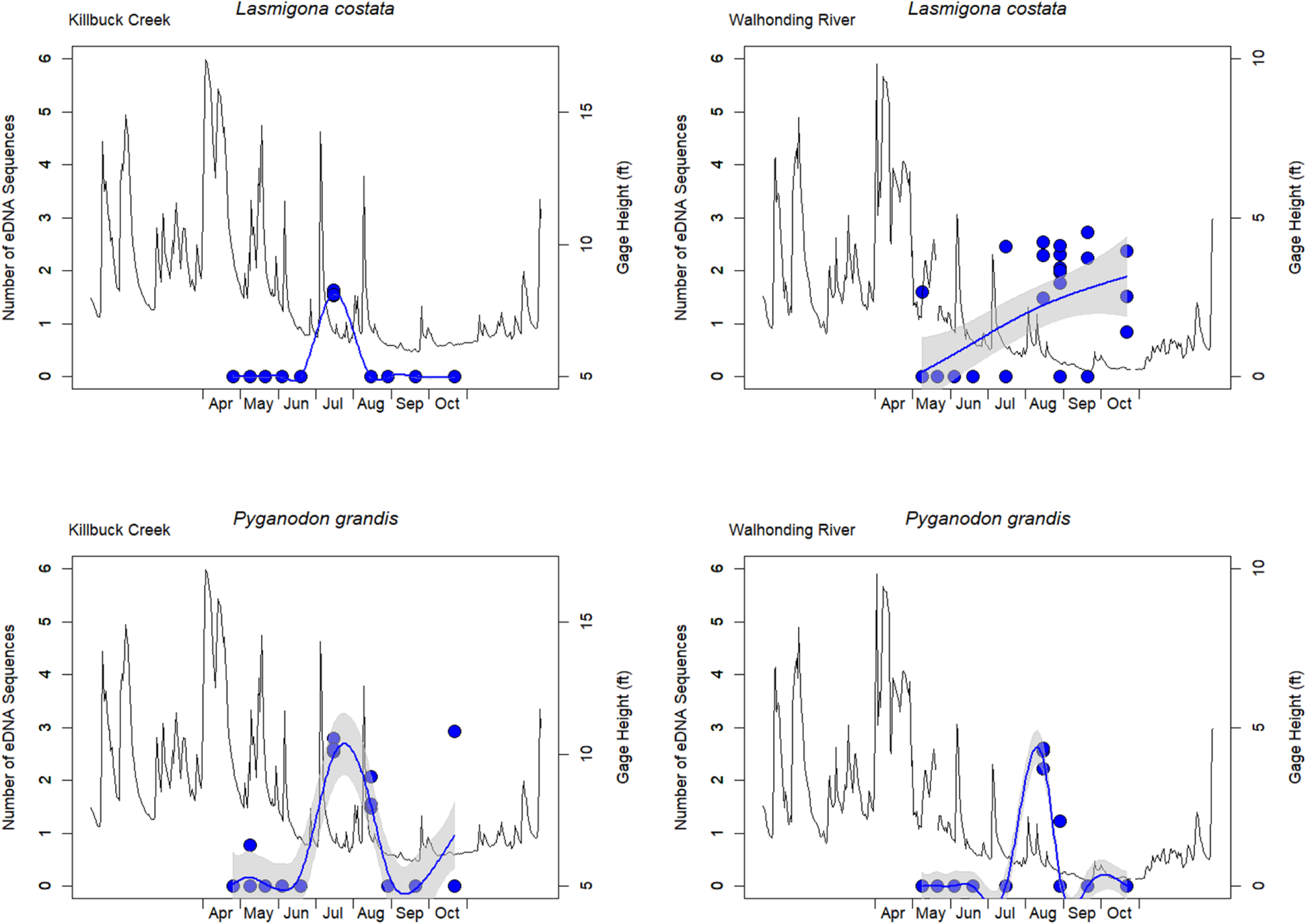
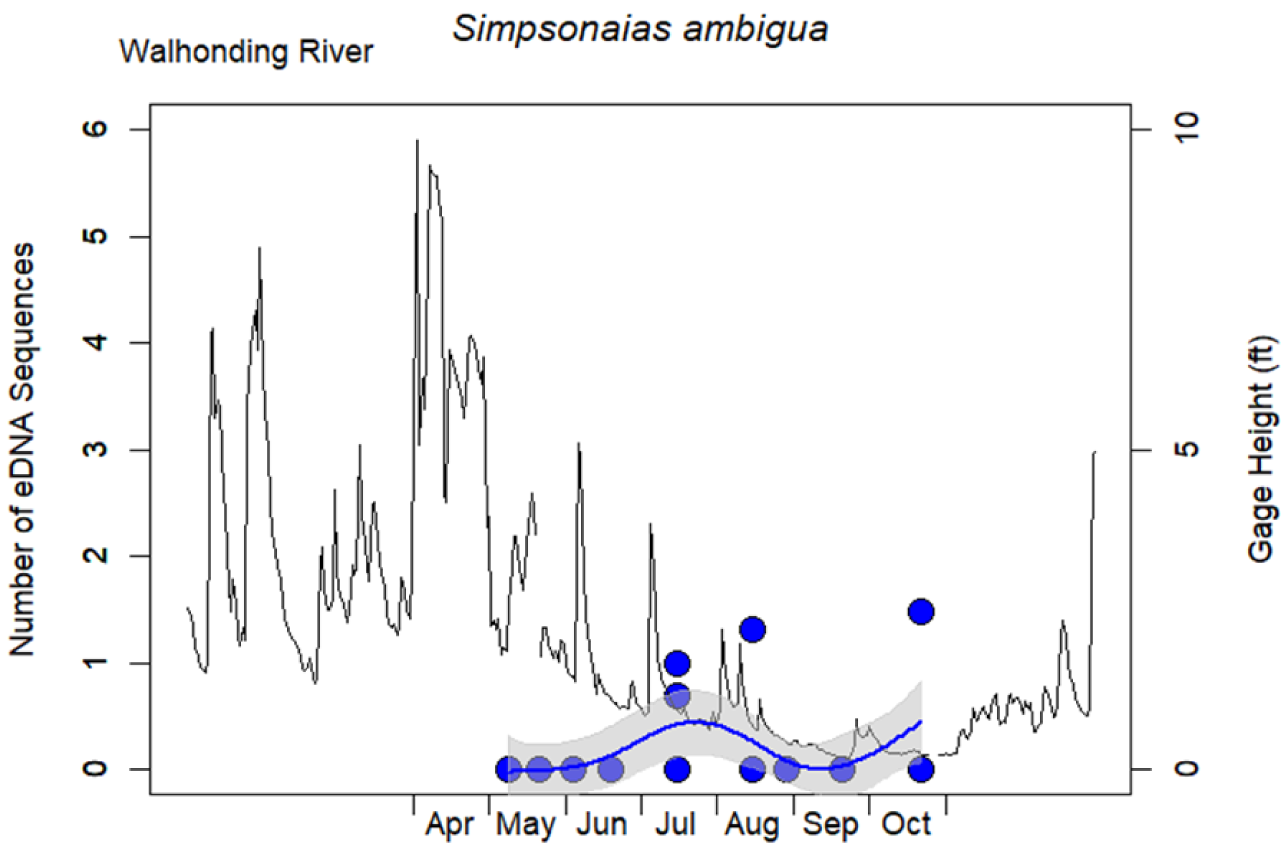
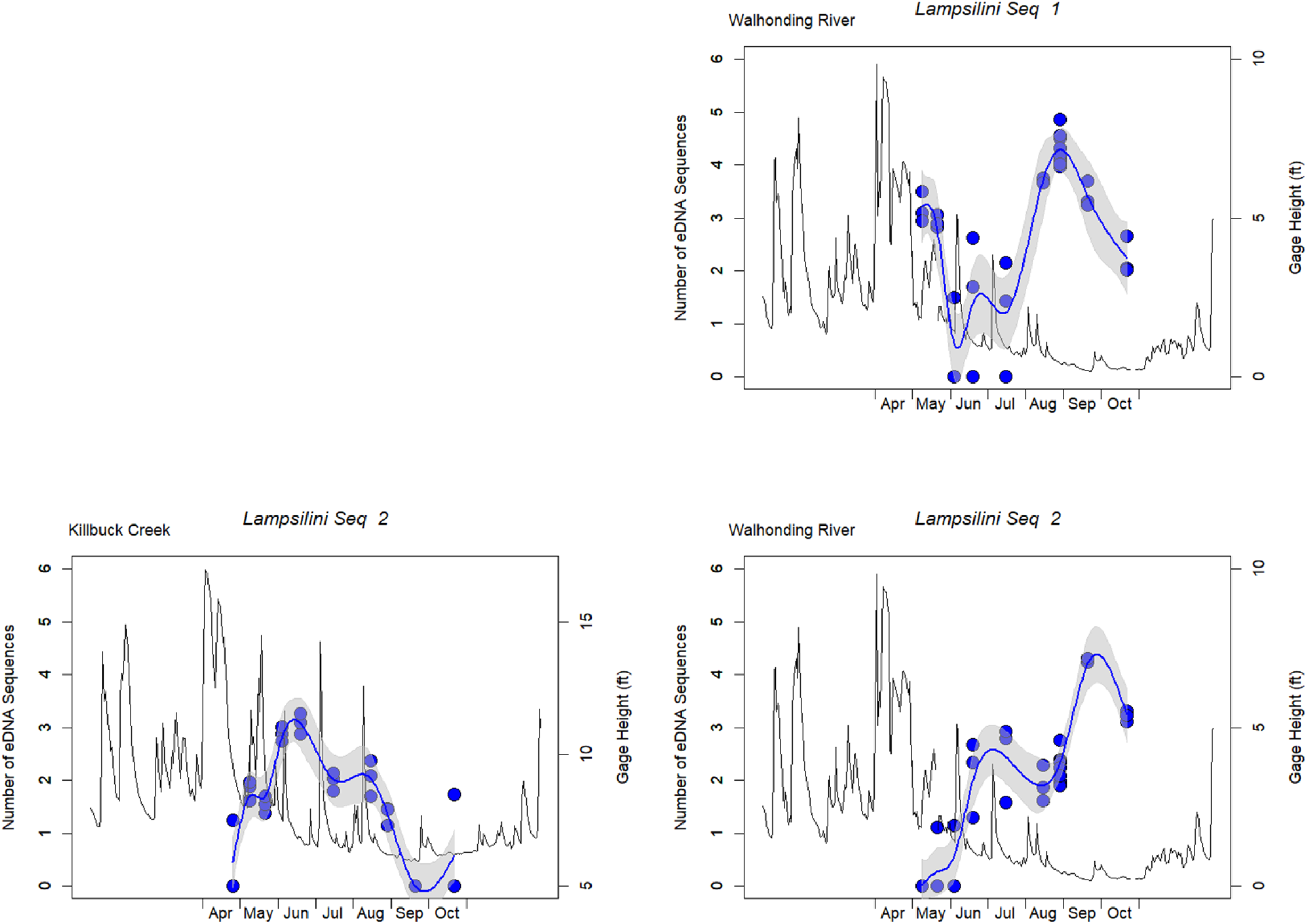
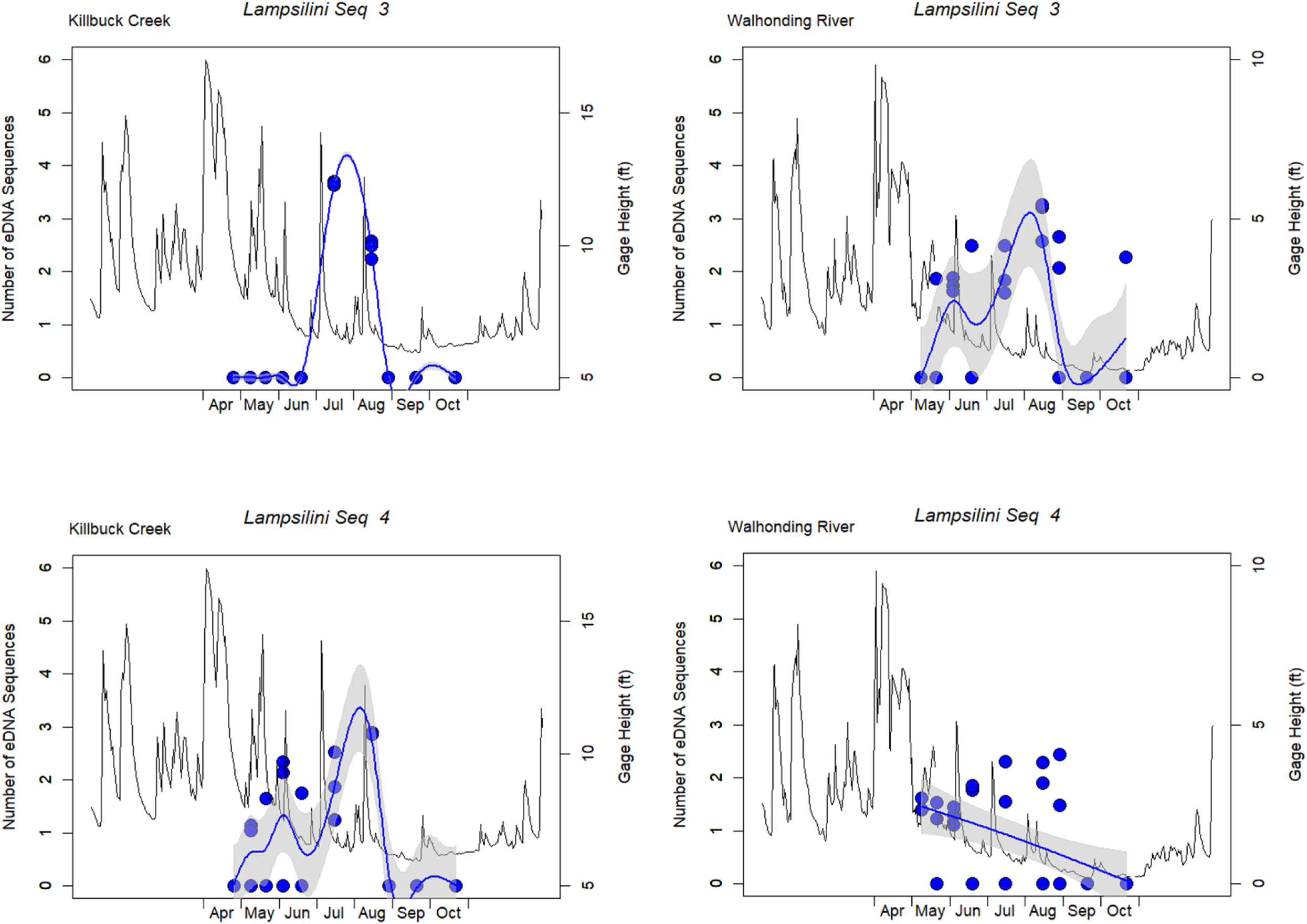
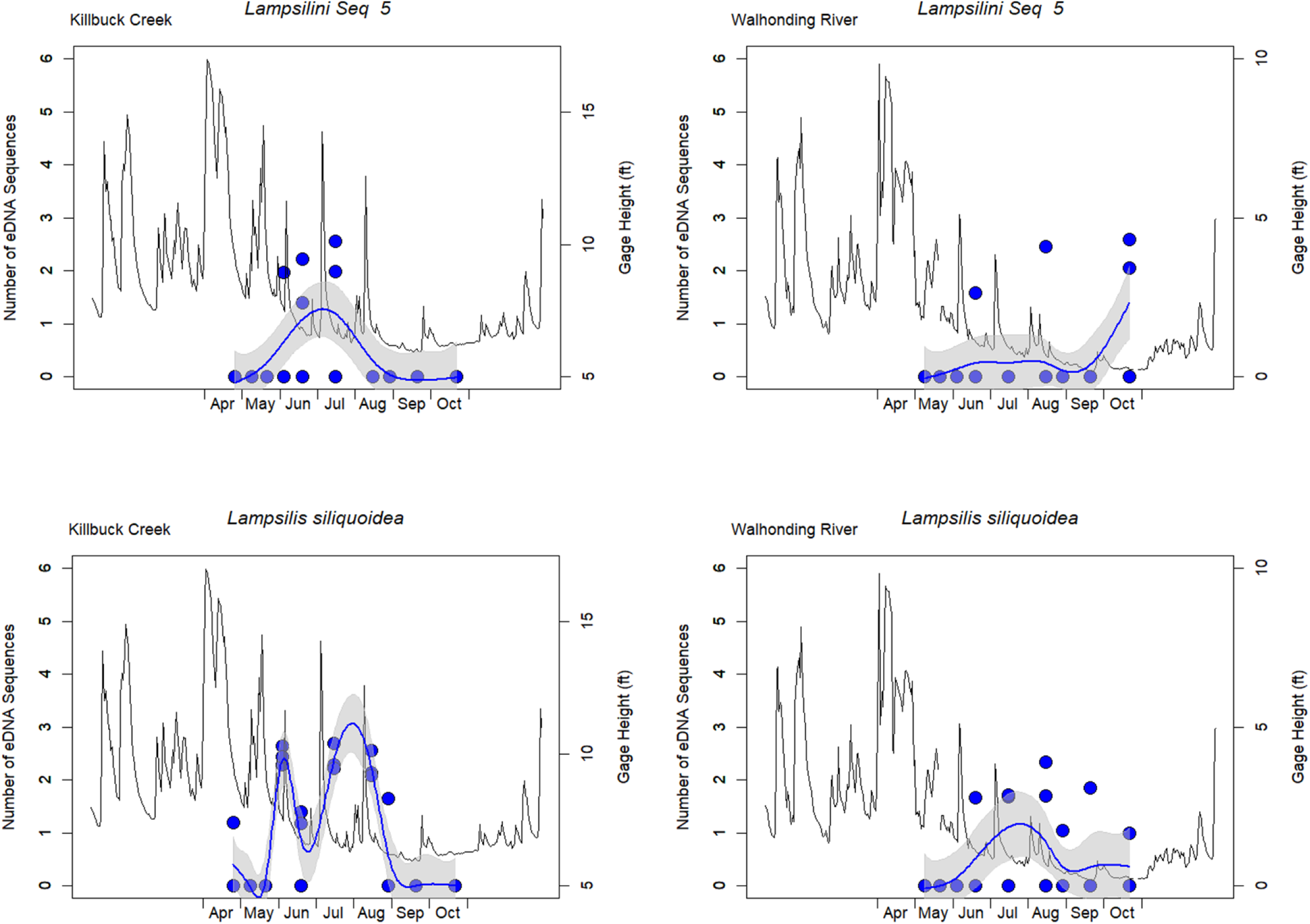
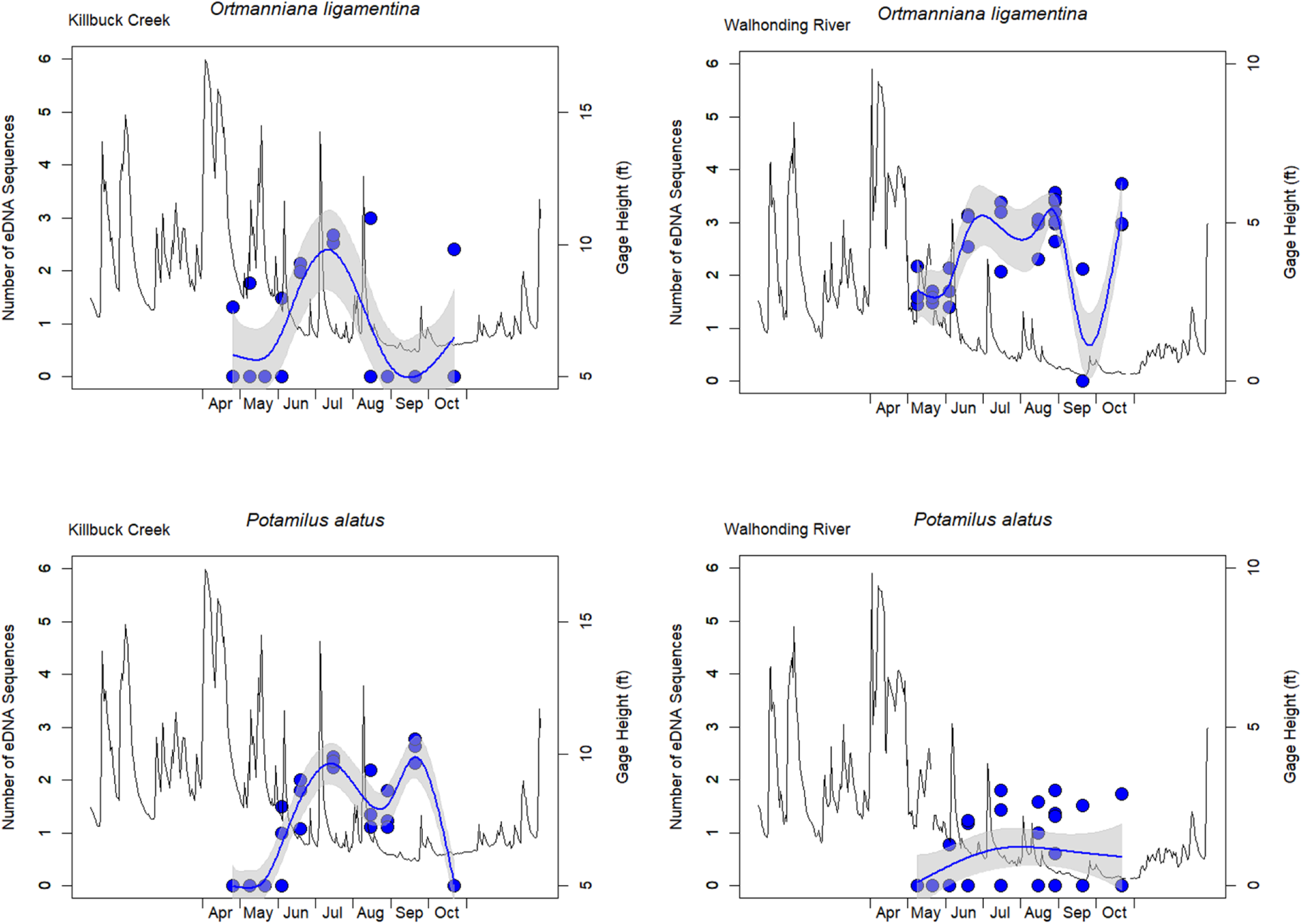
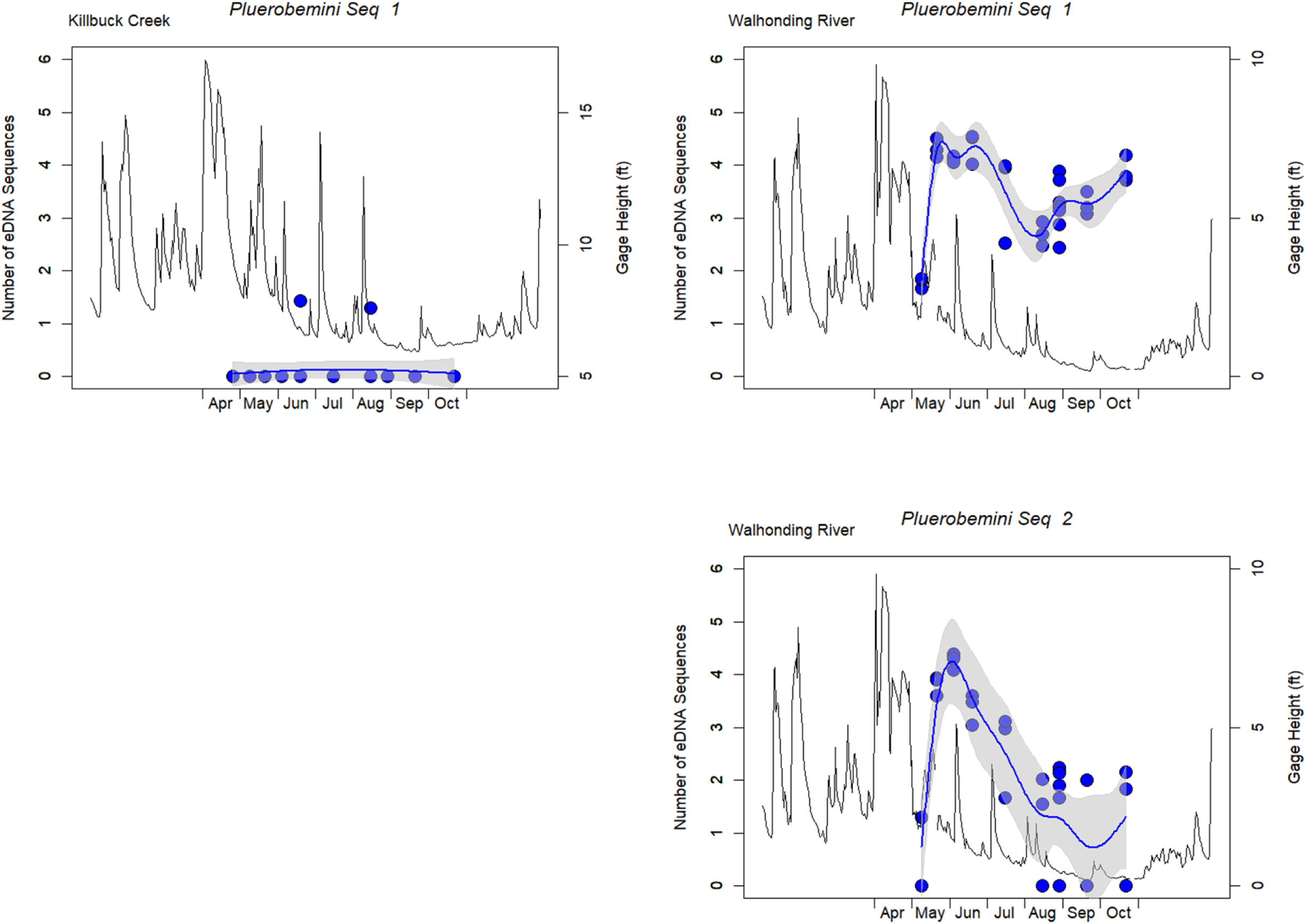
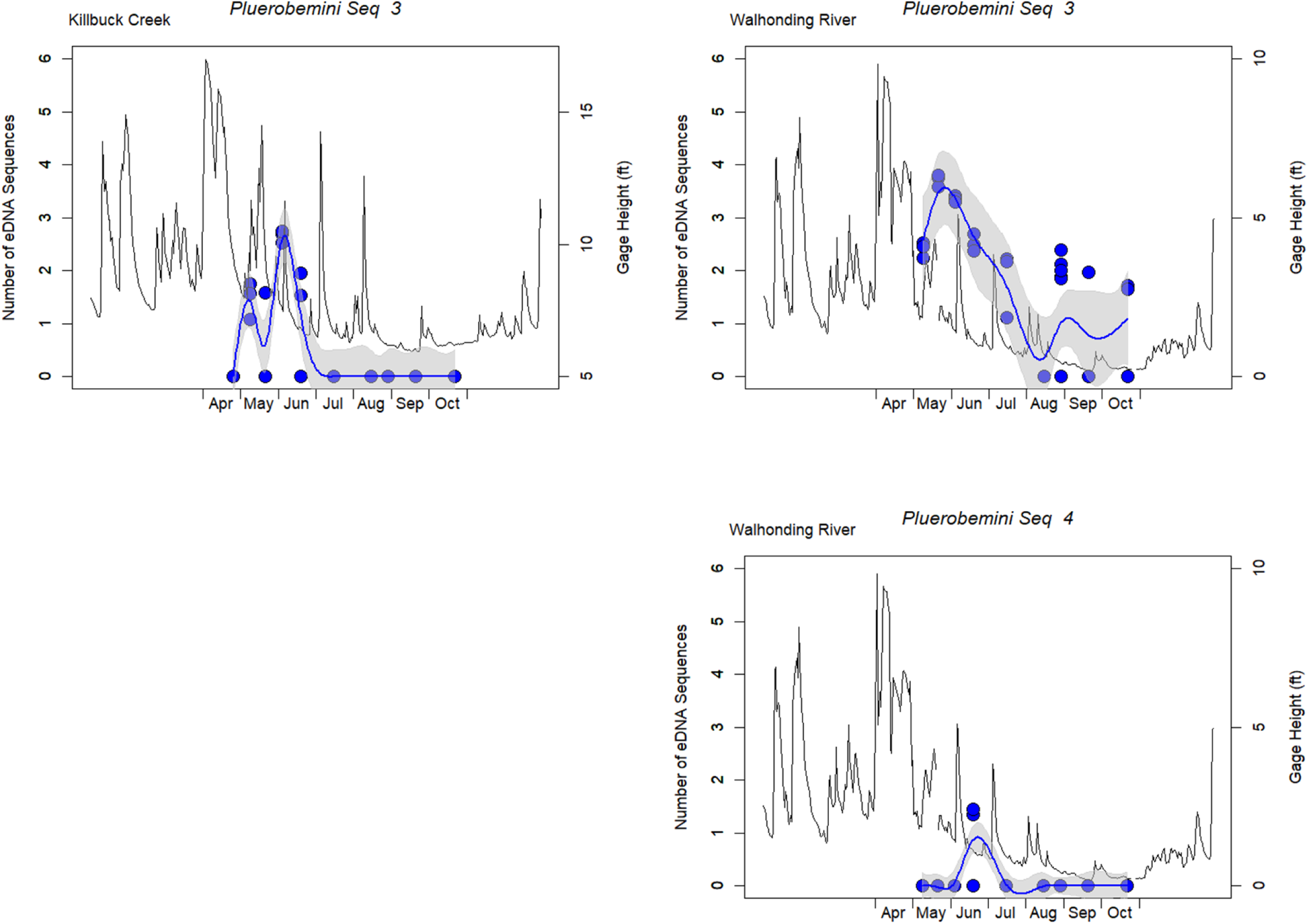
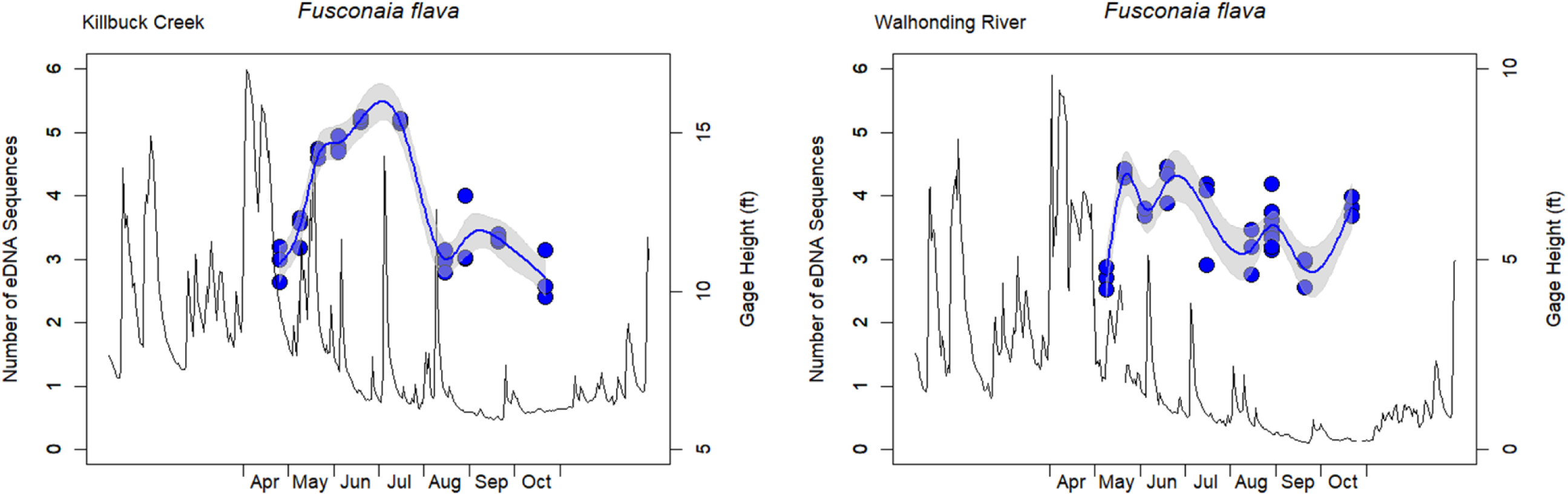
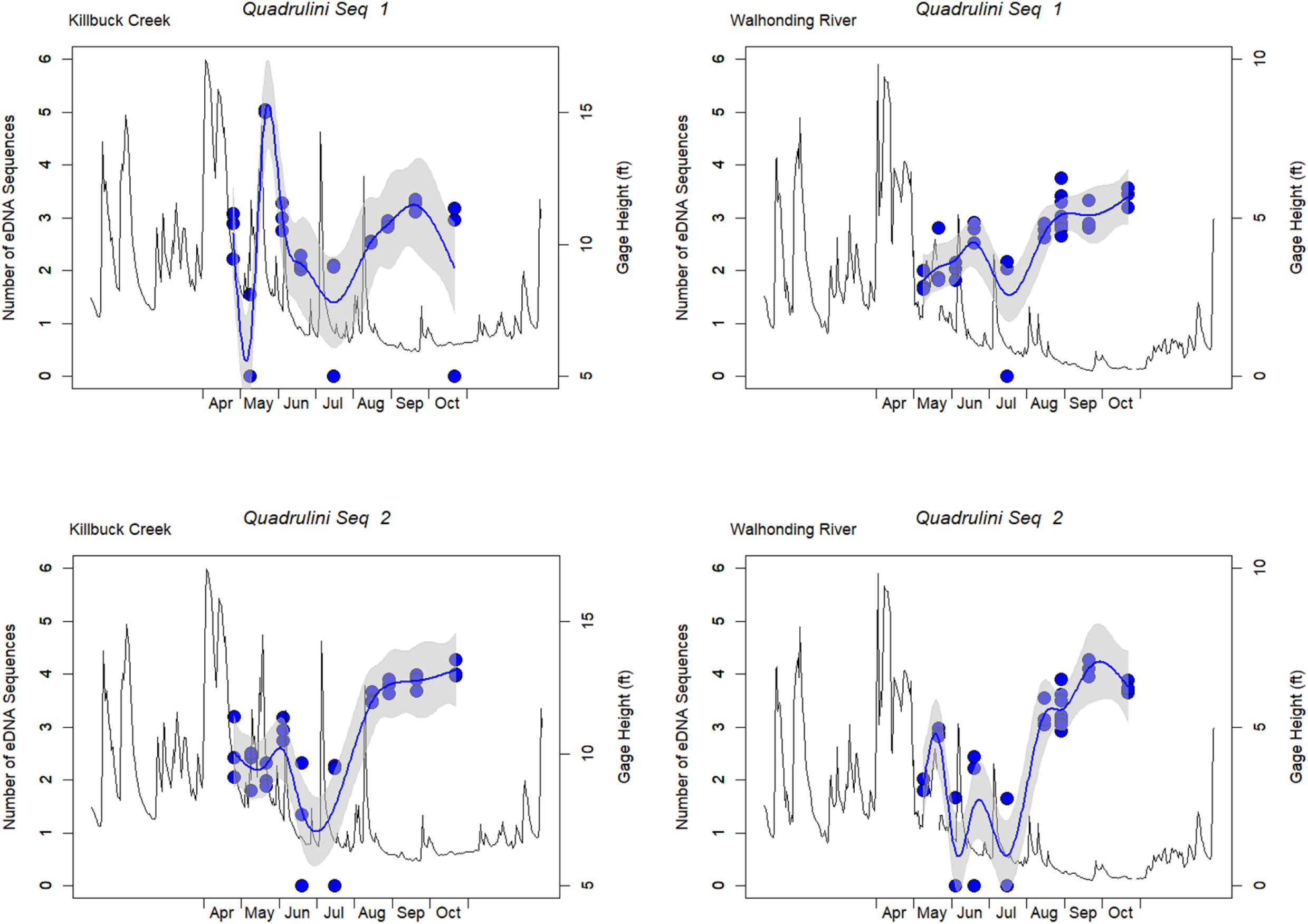
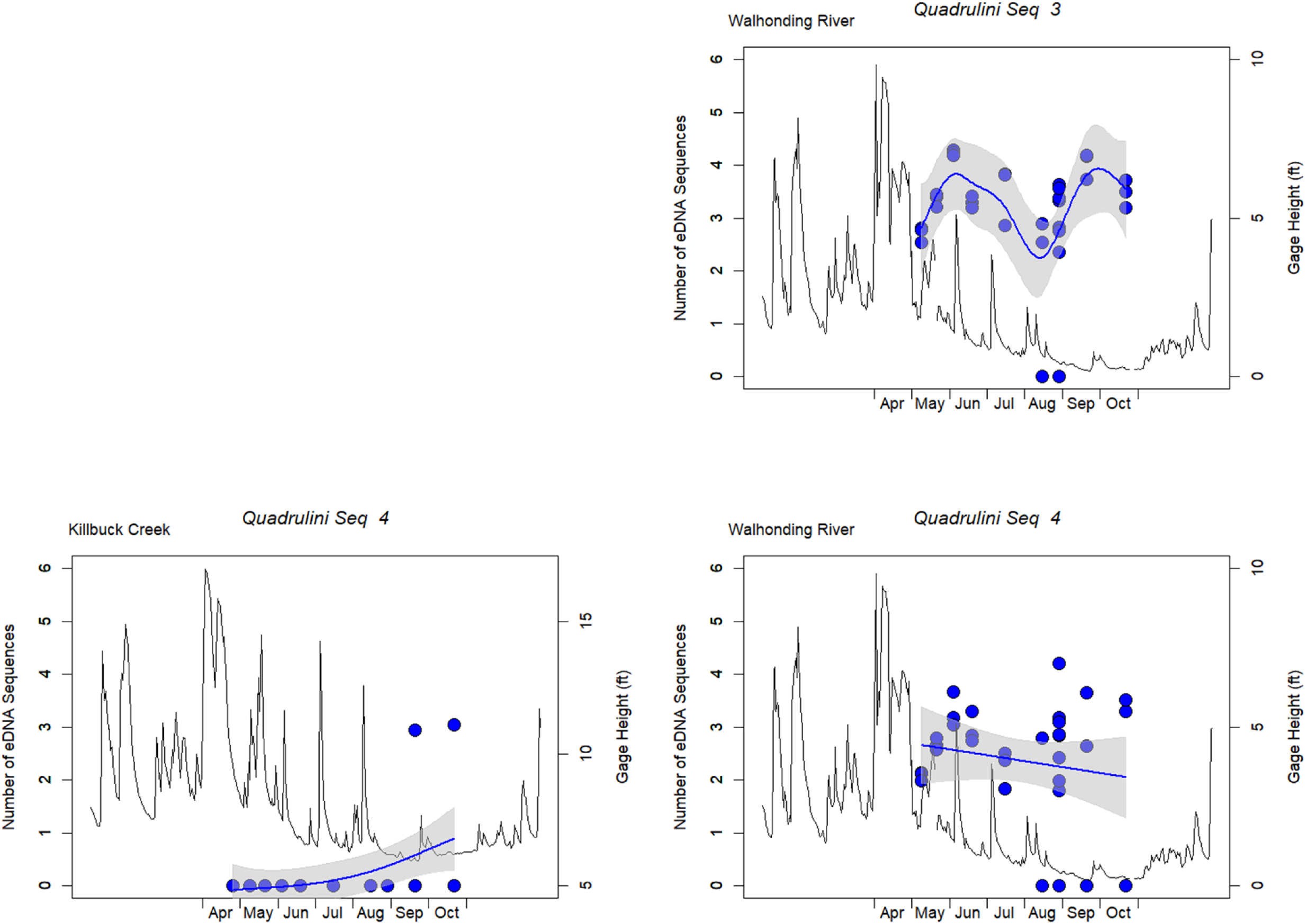
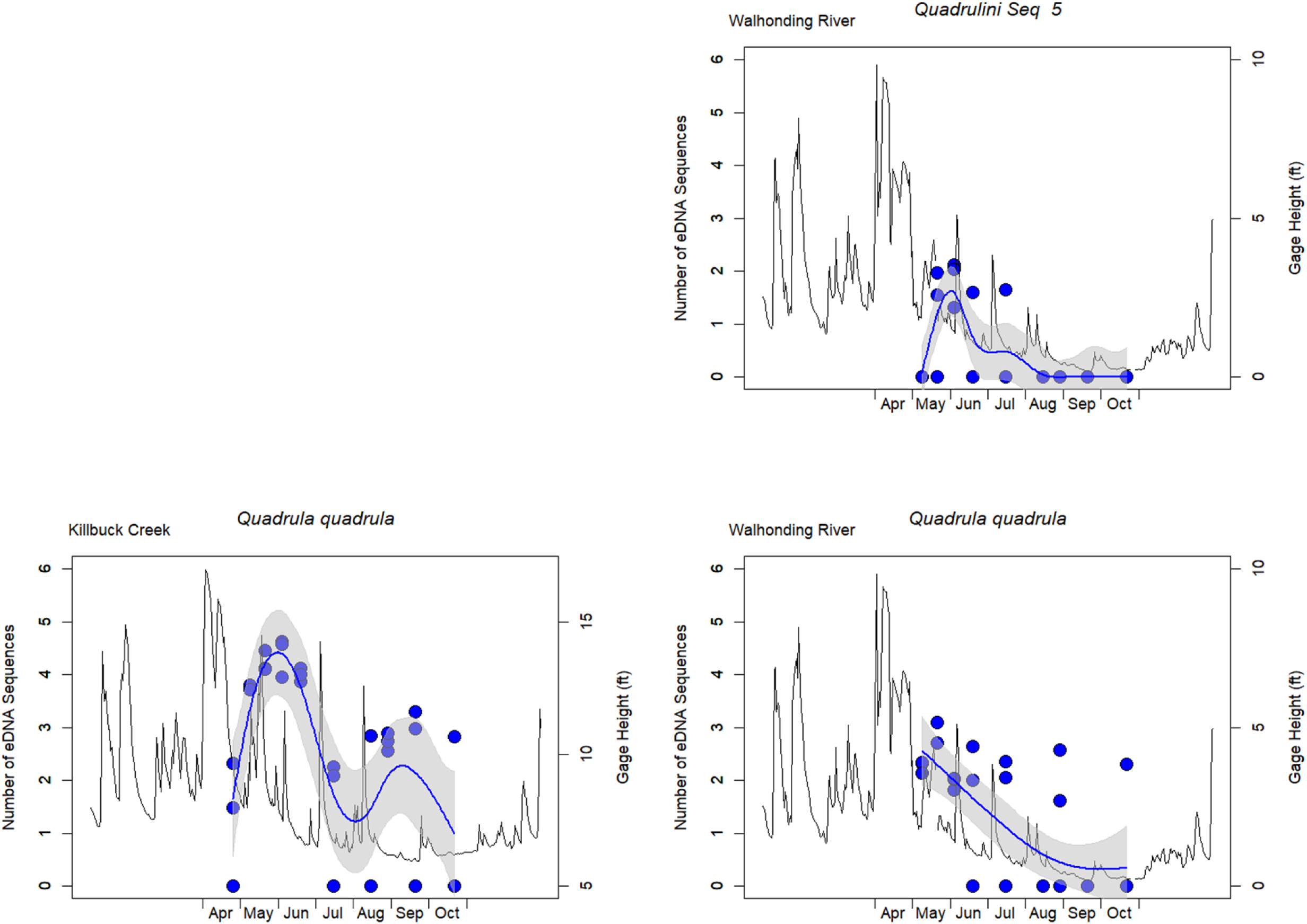
The eDNA abundance (i.e., the number of sequence reads) across all replicates on each sampling date from Killbuck Creek and Walhonding River for each male mitotype MOTU. Plots are organized by species and grouped within the major mussel tribes. The thin black line displays the USGS gage height (ft) across the sampling season.

**Supplementary Figure 3.**
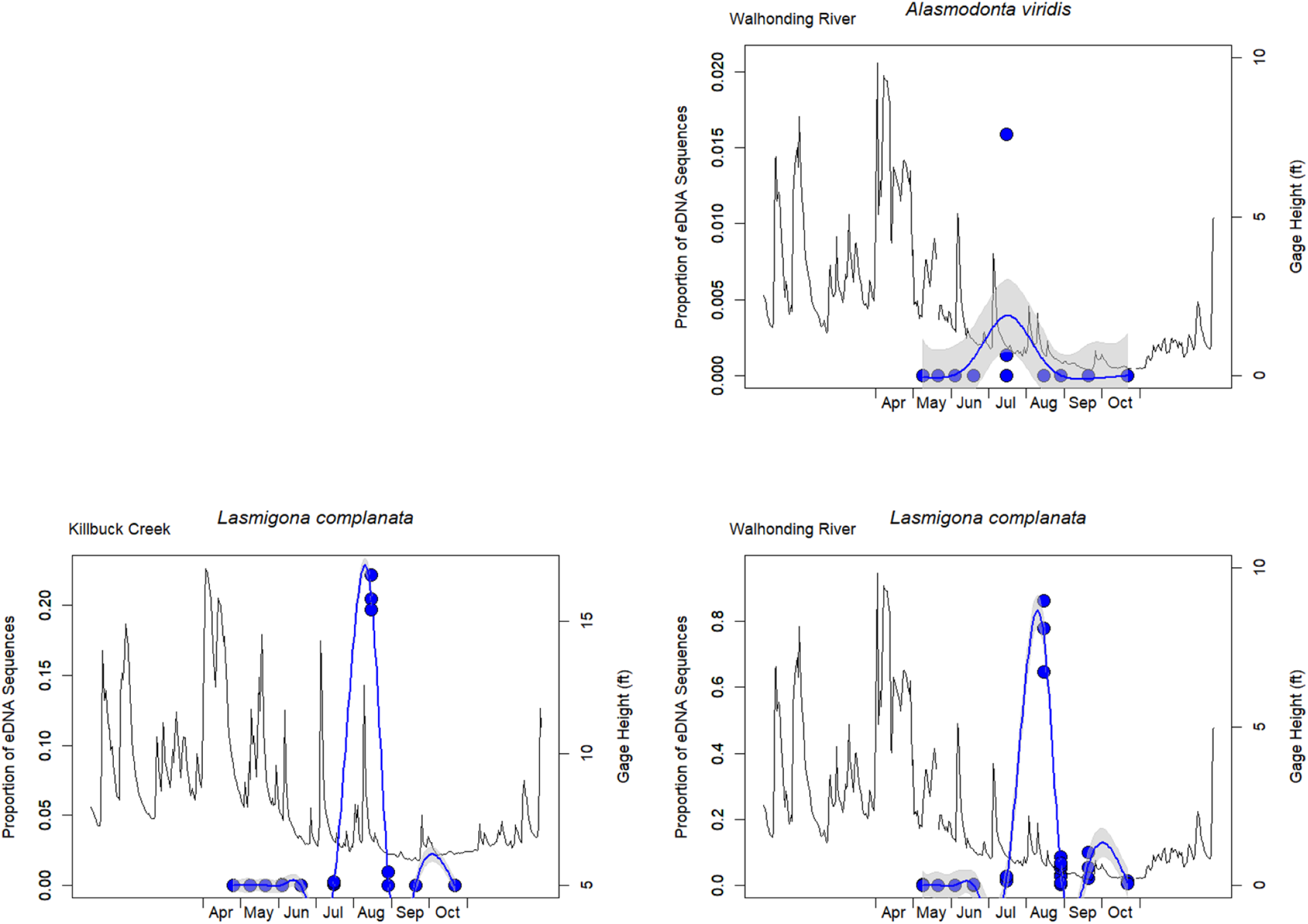
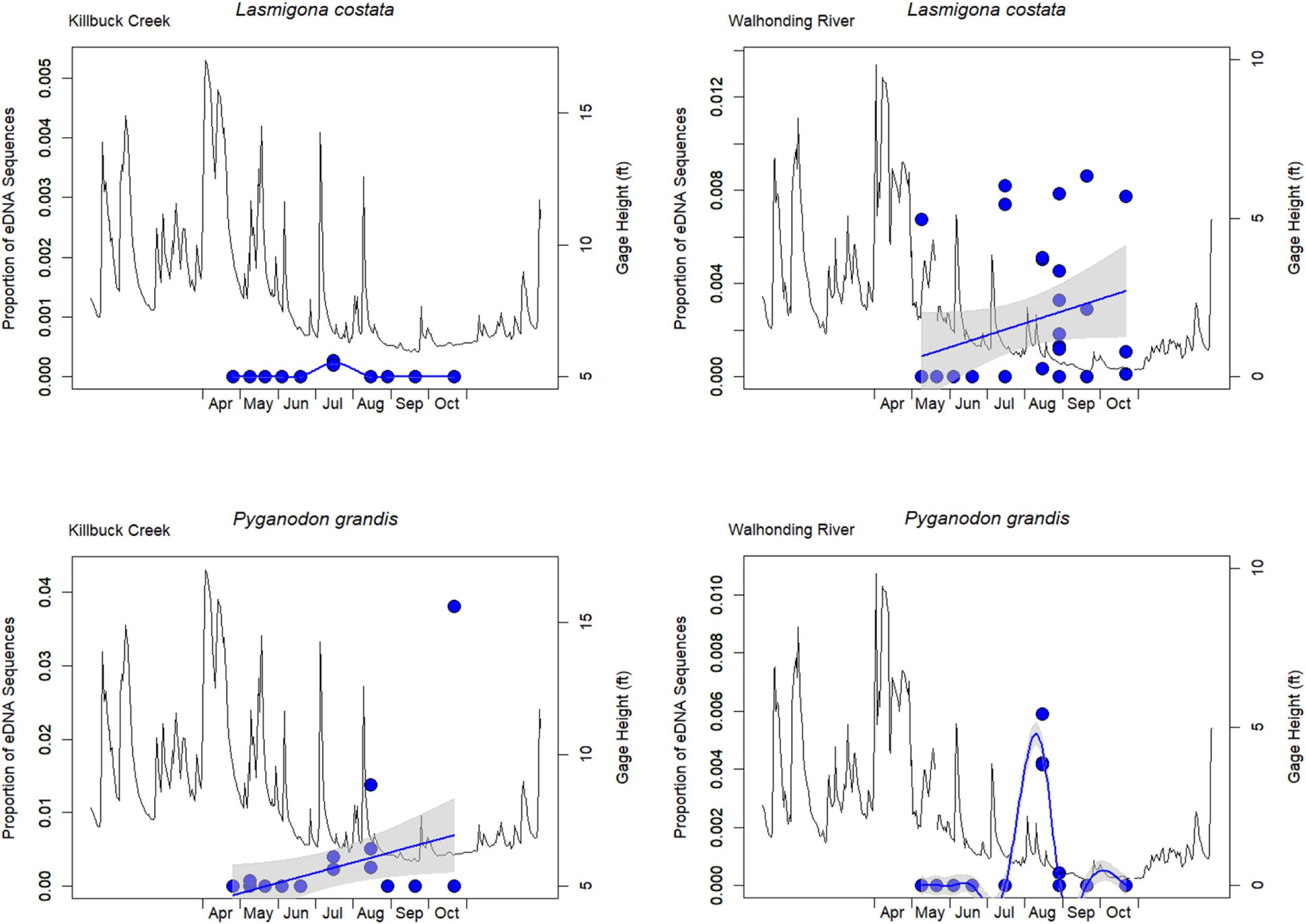
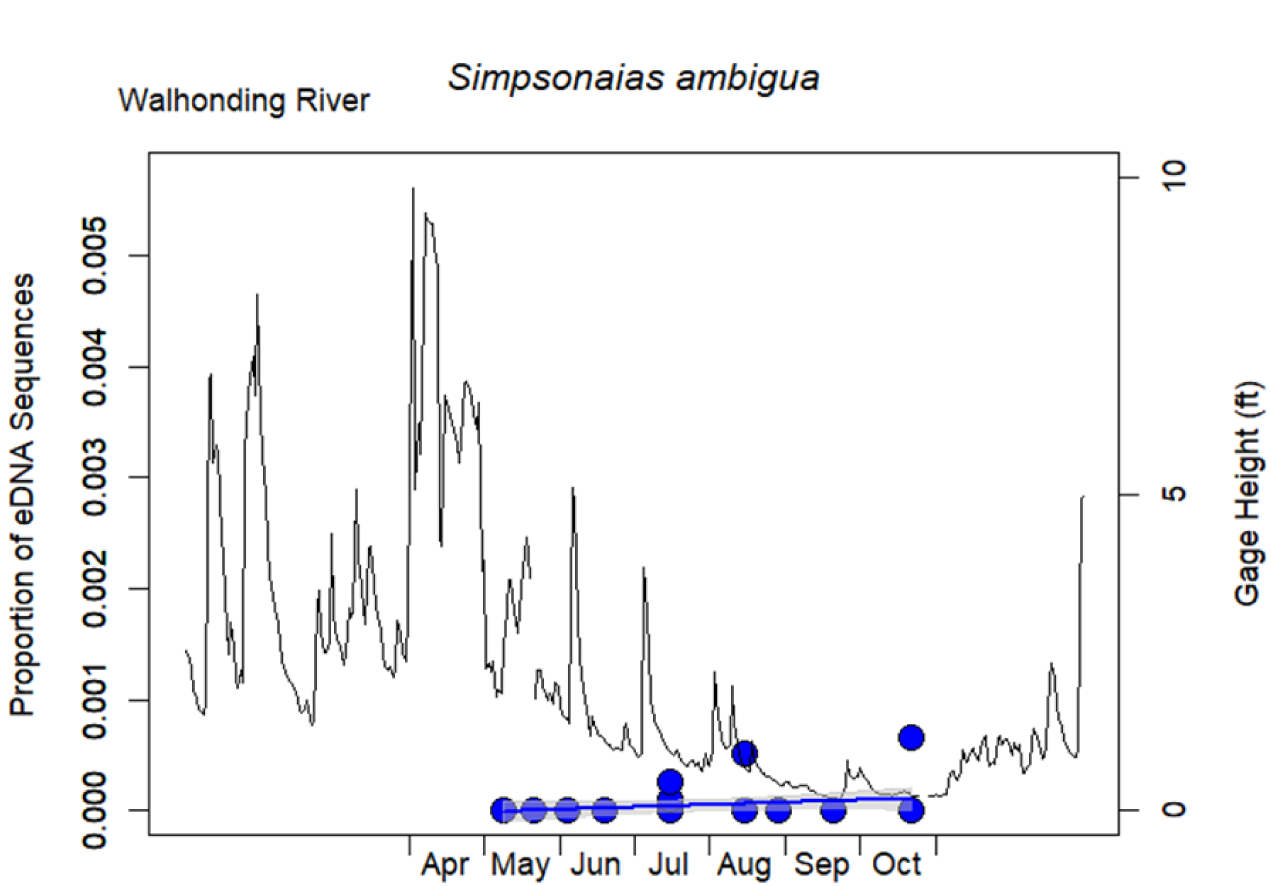
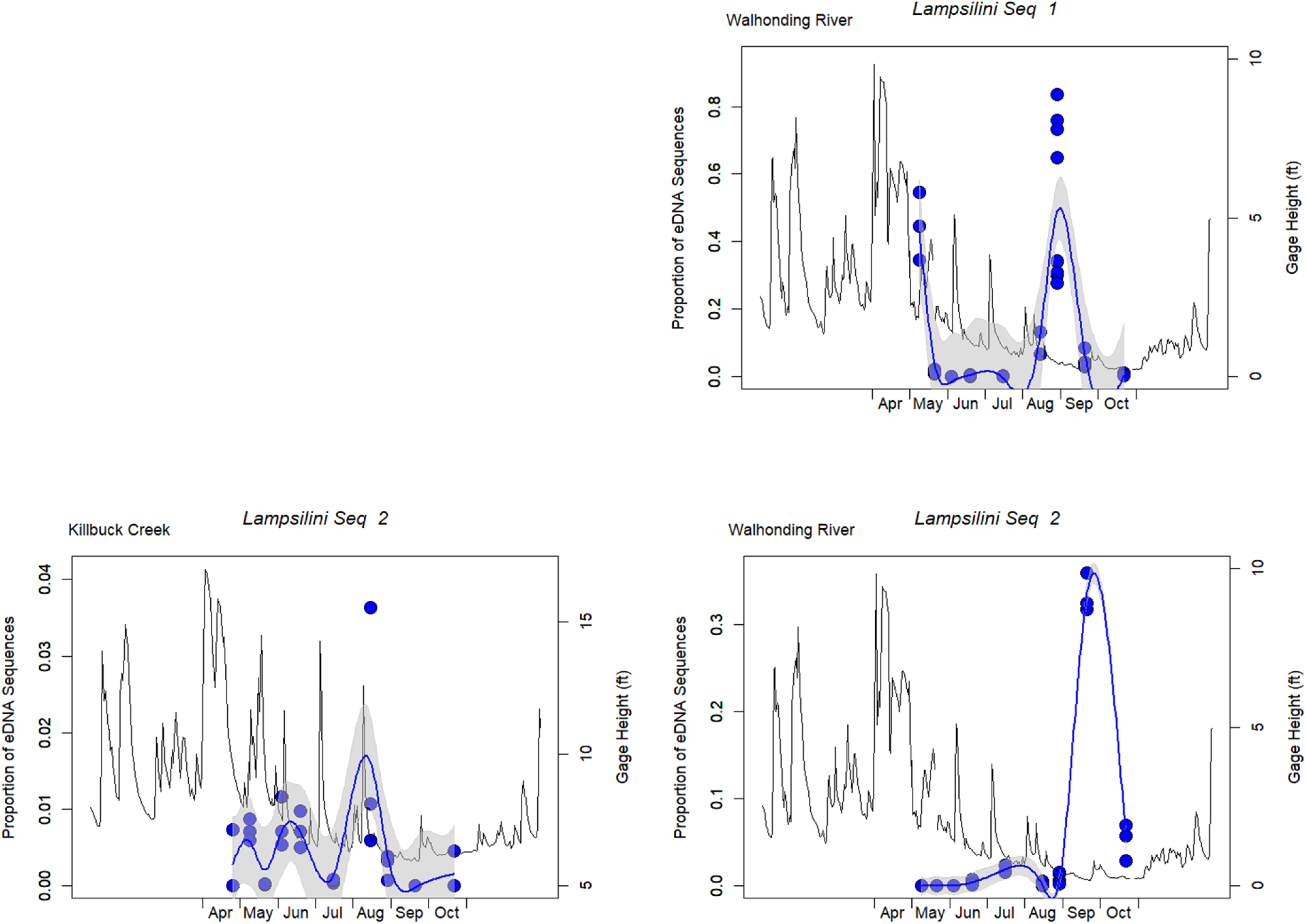
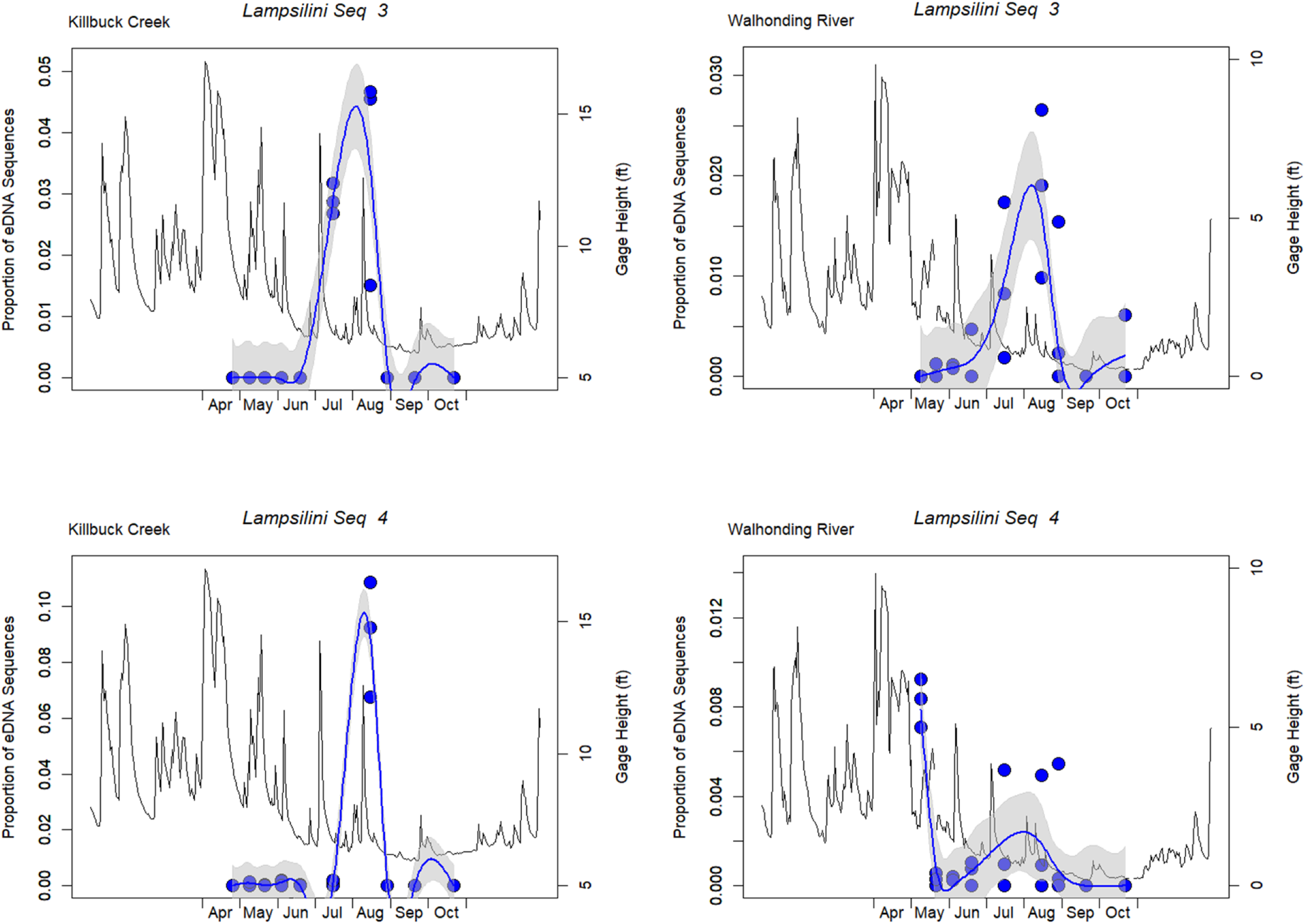
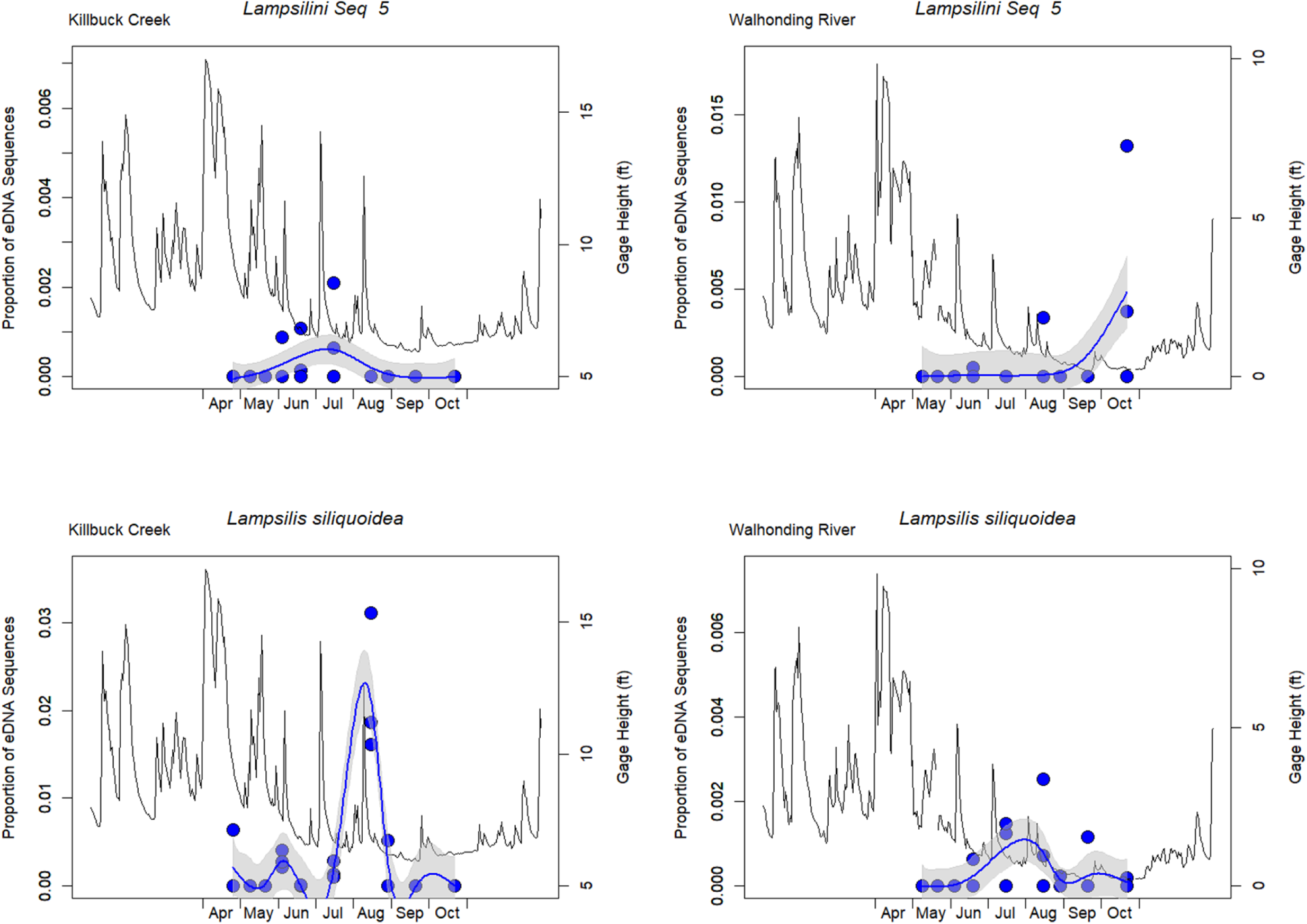
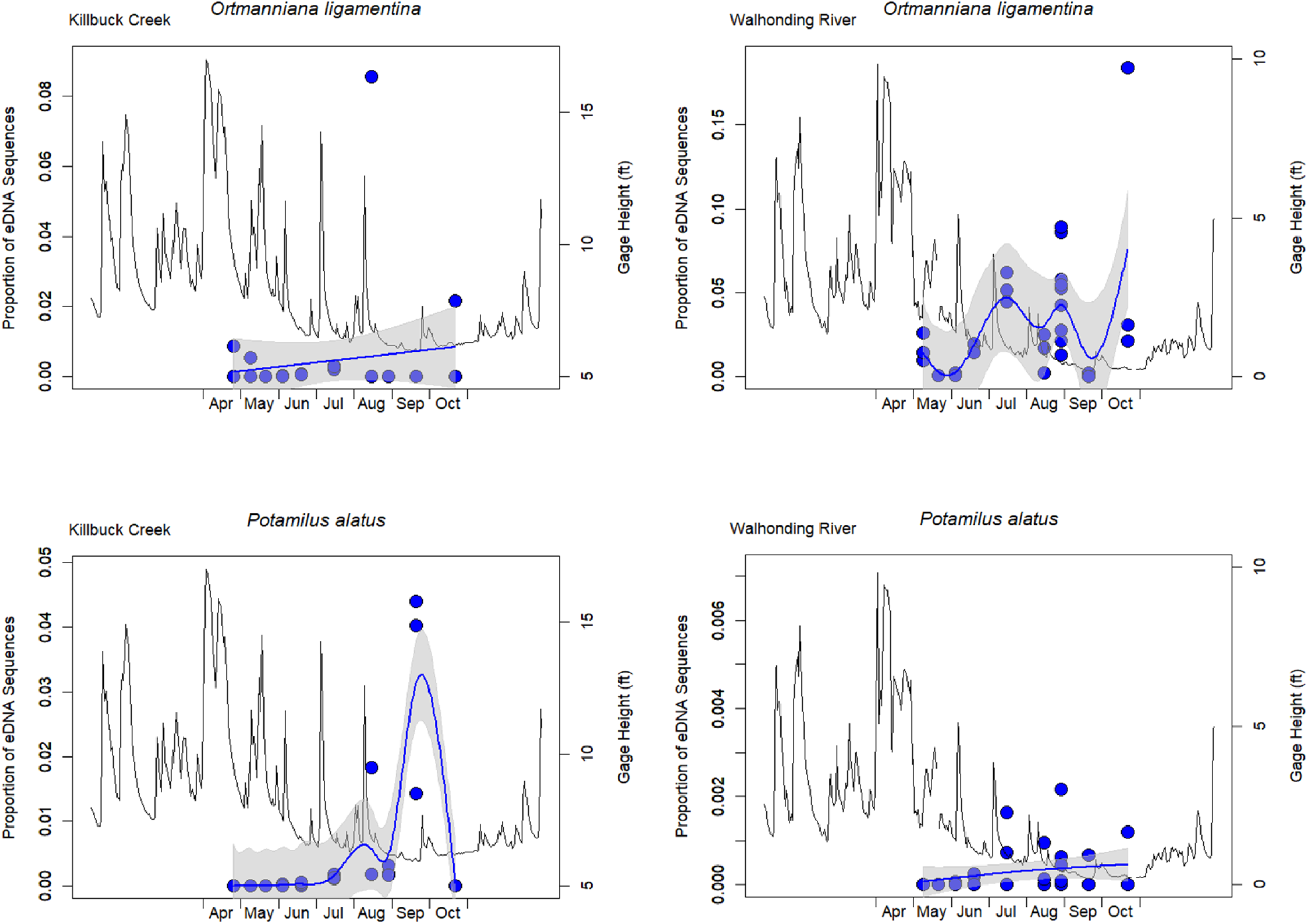
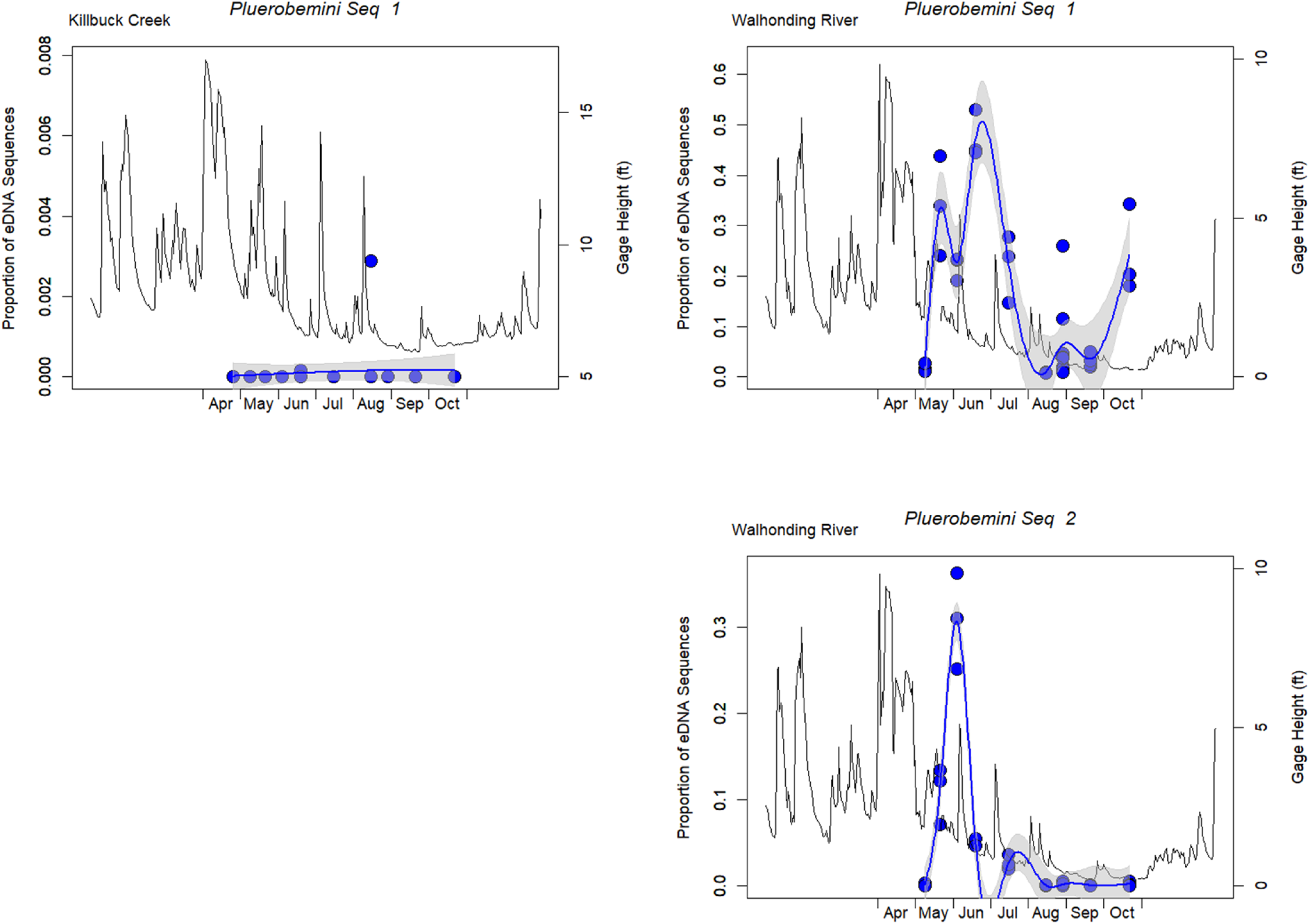
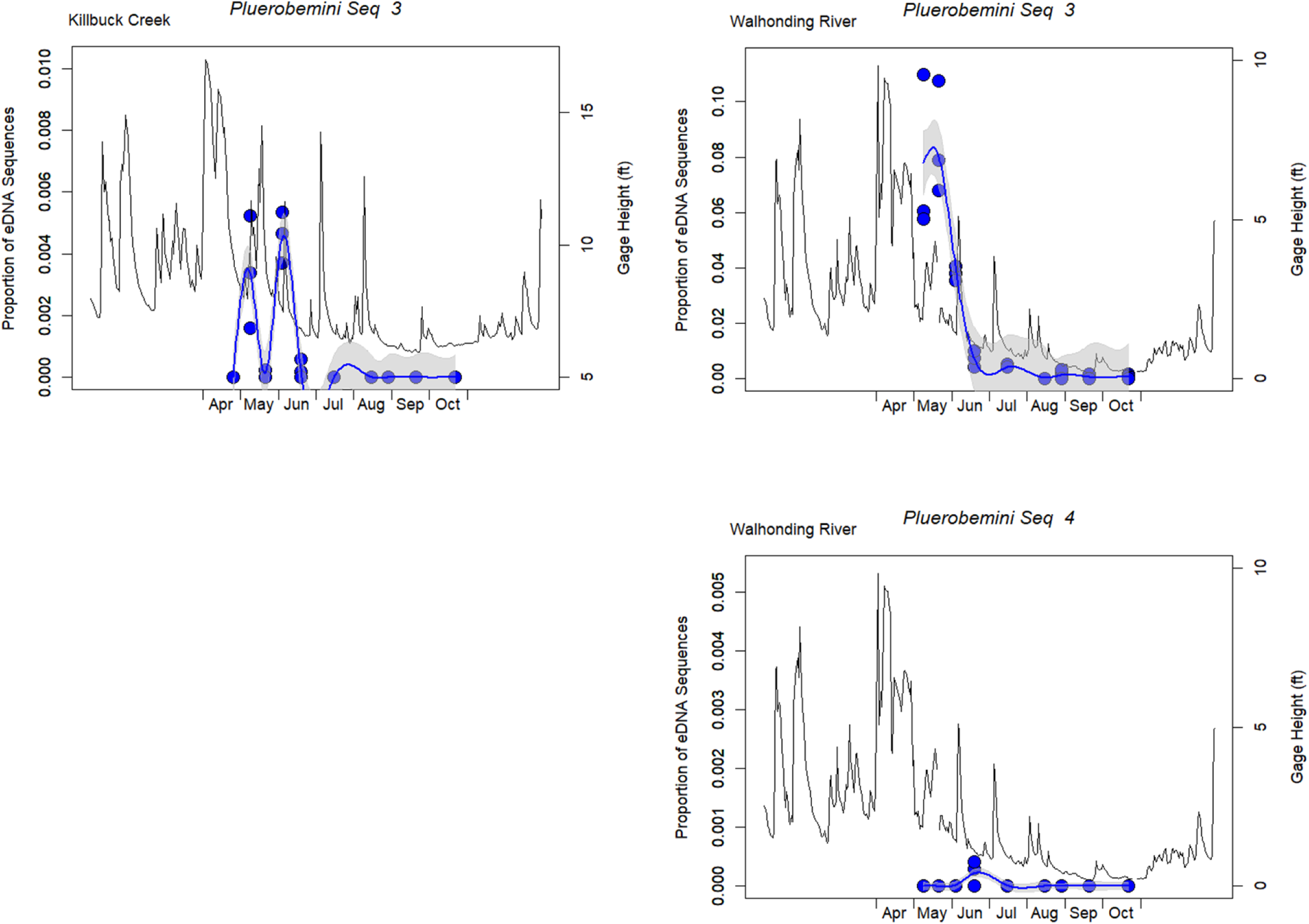
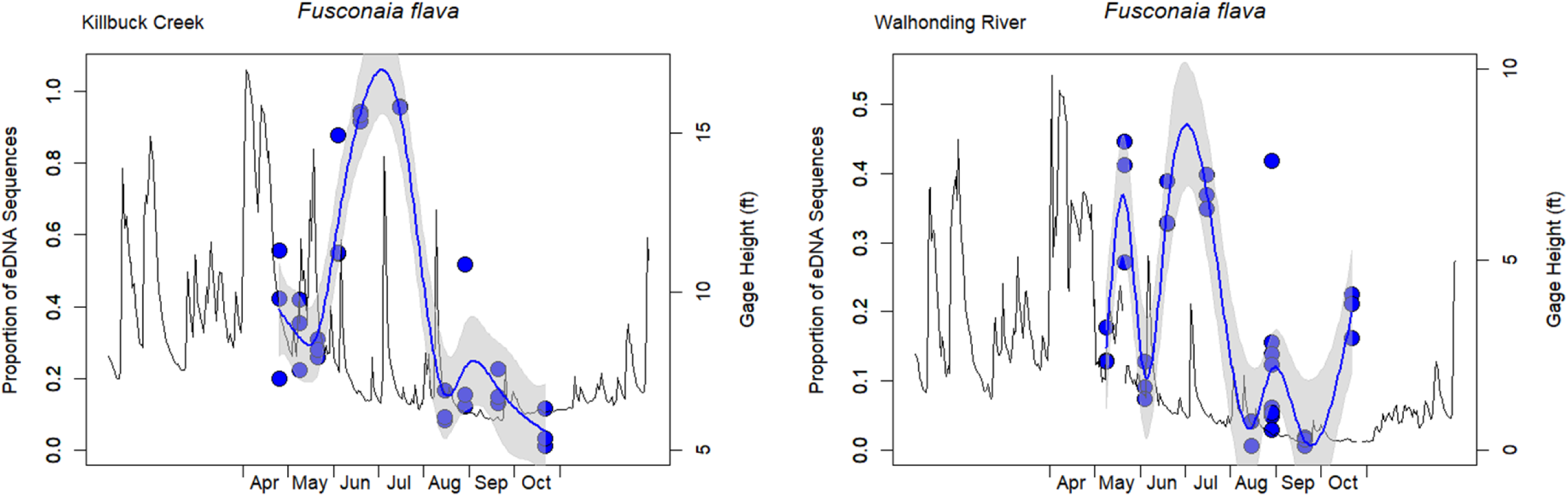
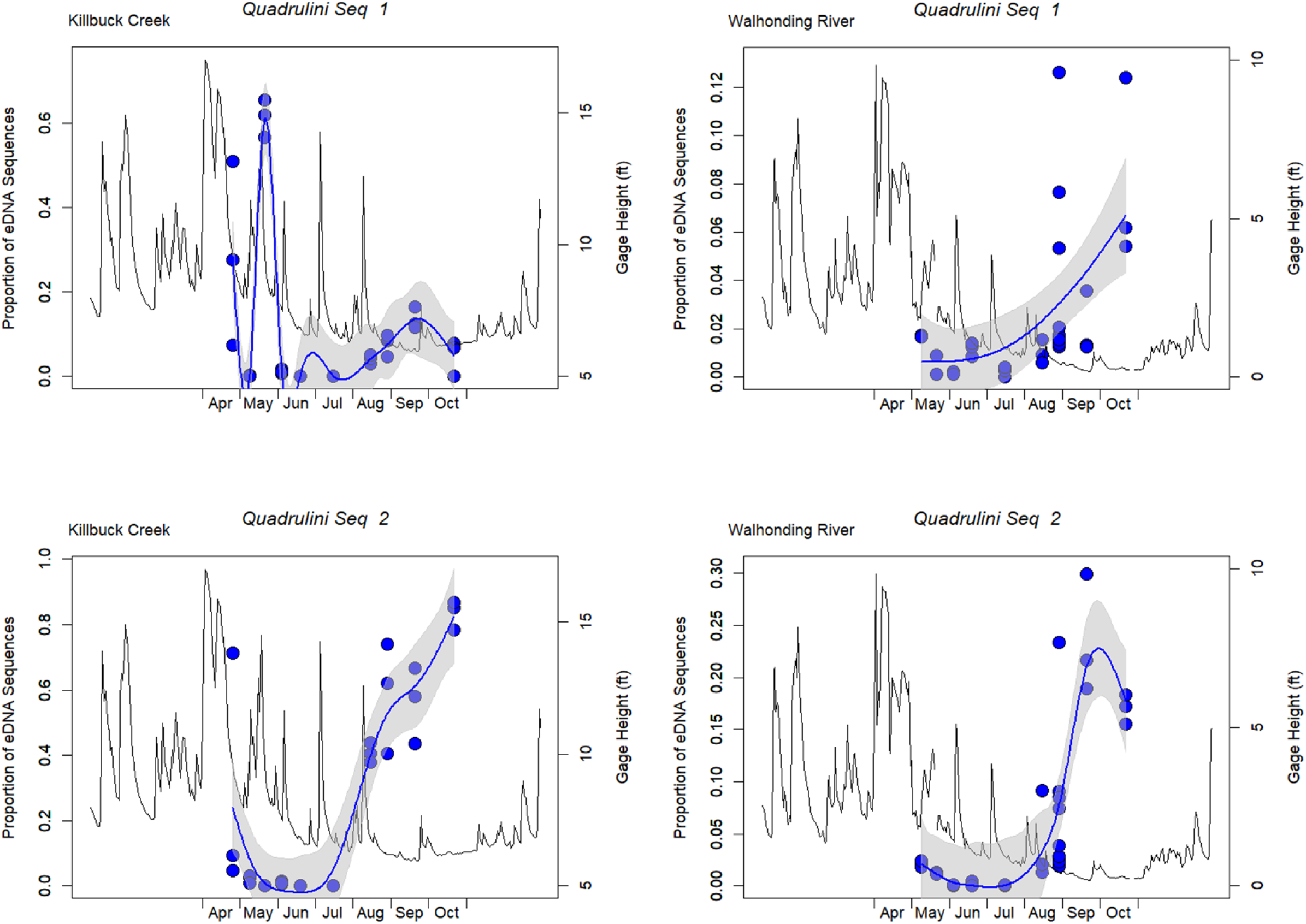
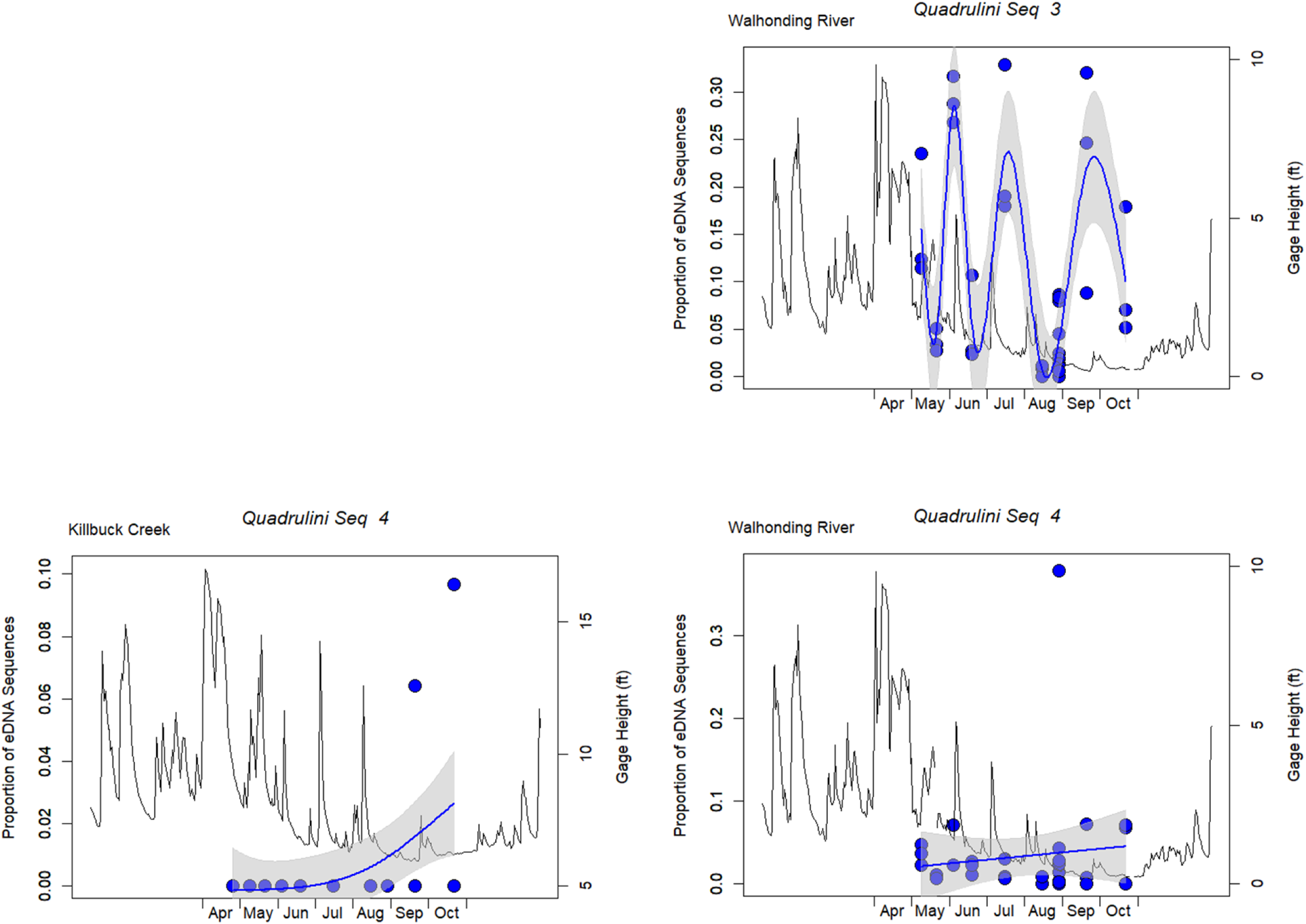
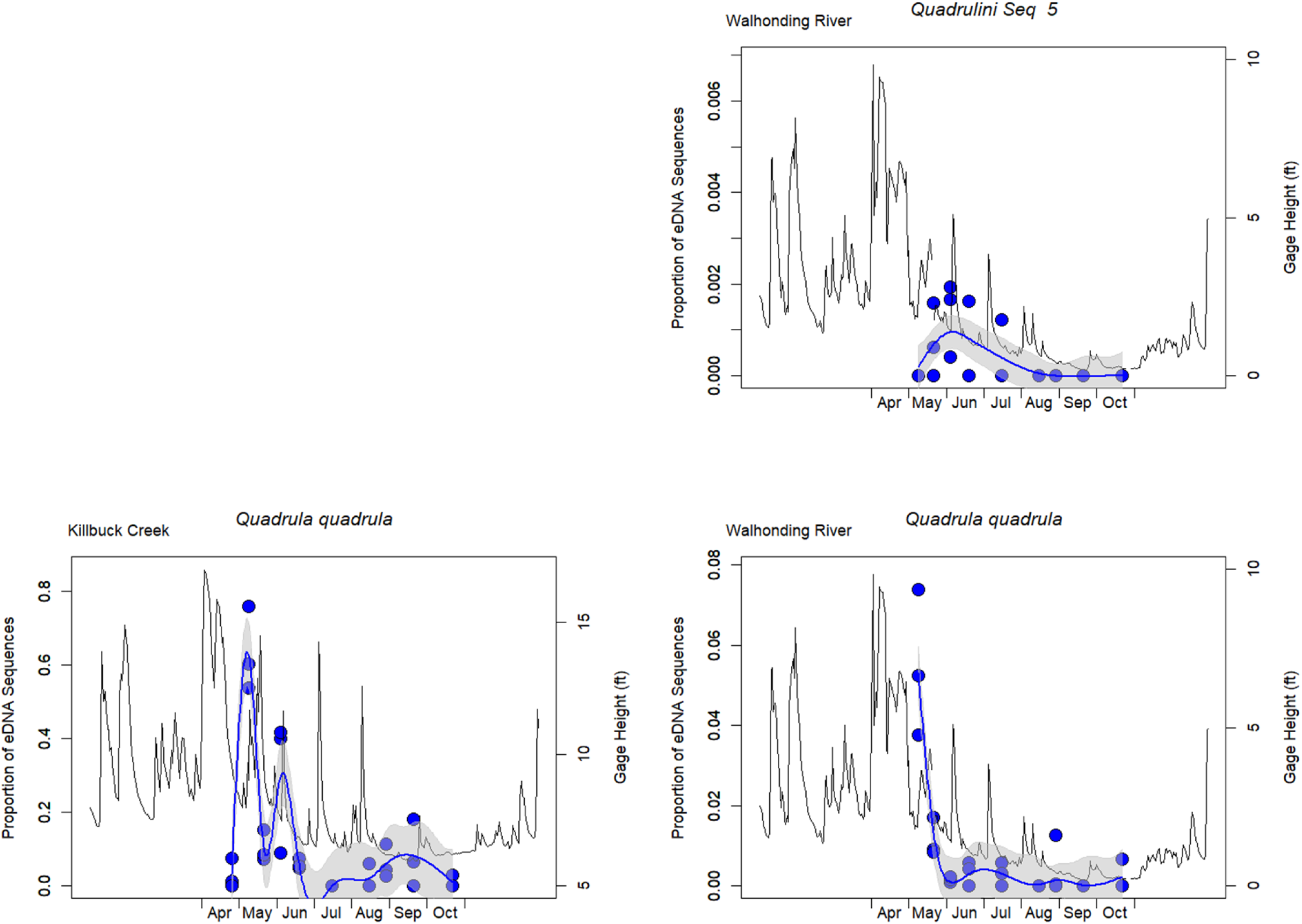
The proportion of the mussel assemblage (i.e., proportion of eDNA sequences) across all replicates on each sampling date from Killbuck Creek and Walhonding River for each male mitotype MOTU. Plots are organized by species and grouped within the major mussel tribes. The thin black line displays the USGS gage height (ft) across the sampling season.

**Supplementary Figure 4.**
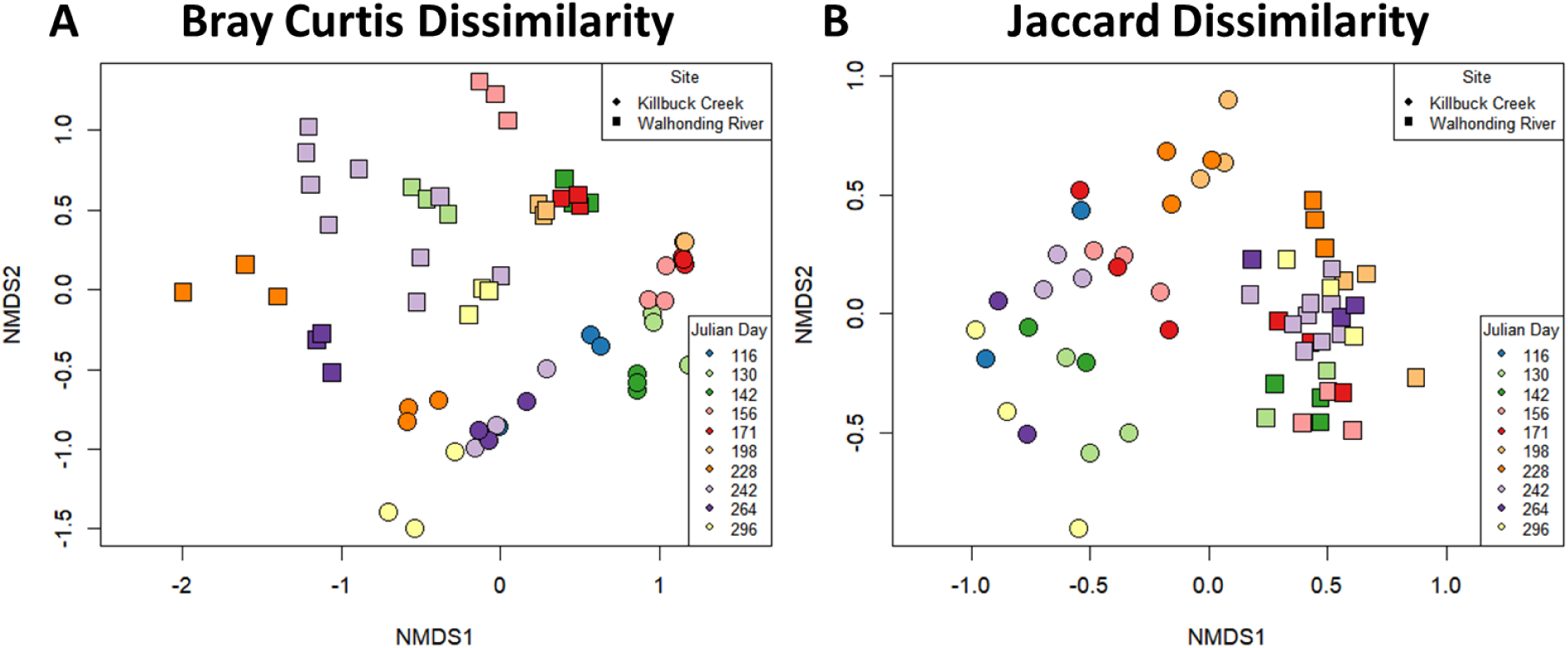
Non-metric multidimensional scaling plots for (A) Bray-Curtis Dissimilarity and (B) Jaccard Dissimilarity for the male mitotype assemblage detected with environmental DNA.

**Supplementary Figure 5.**
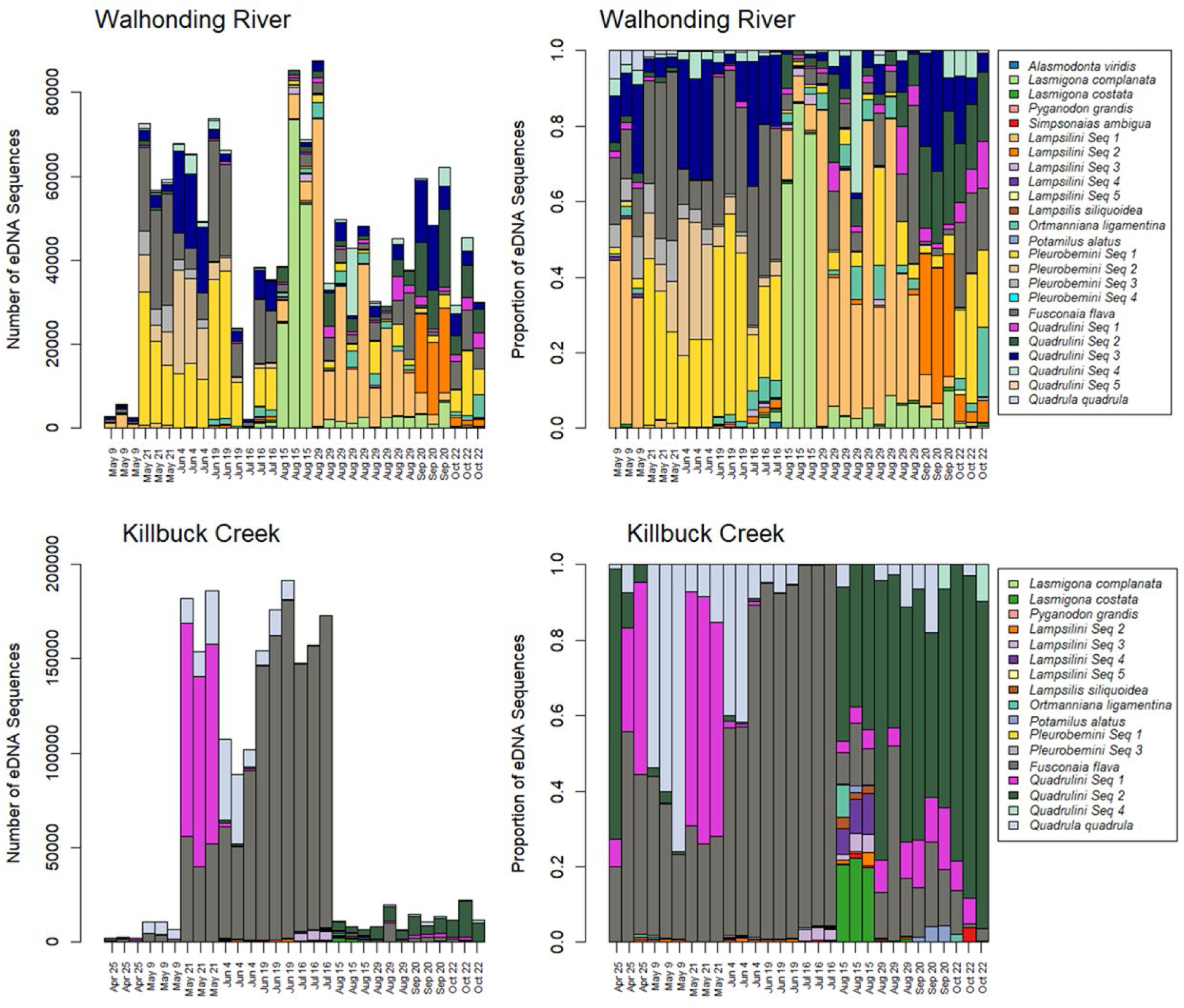
Number of male mitotype environmental DNA sequences and proportion of male mitotype environmental DNA sequences detected across sampling water replicates in Killbuck Creek and Walhonding River.

**Supplementary Figure 6.**
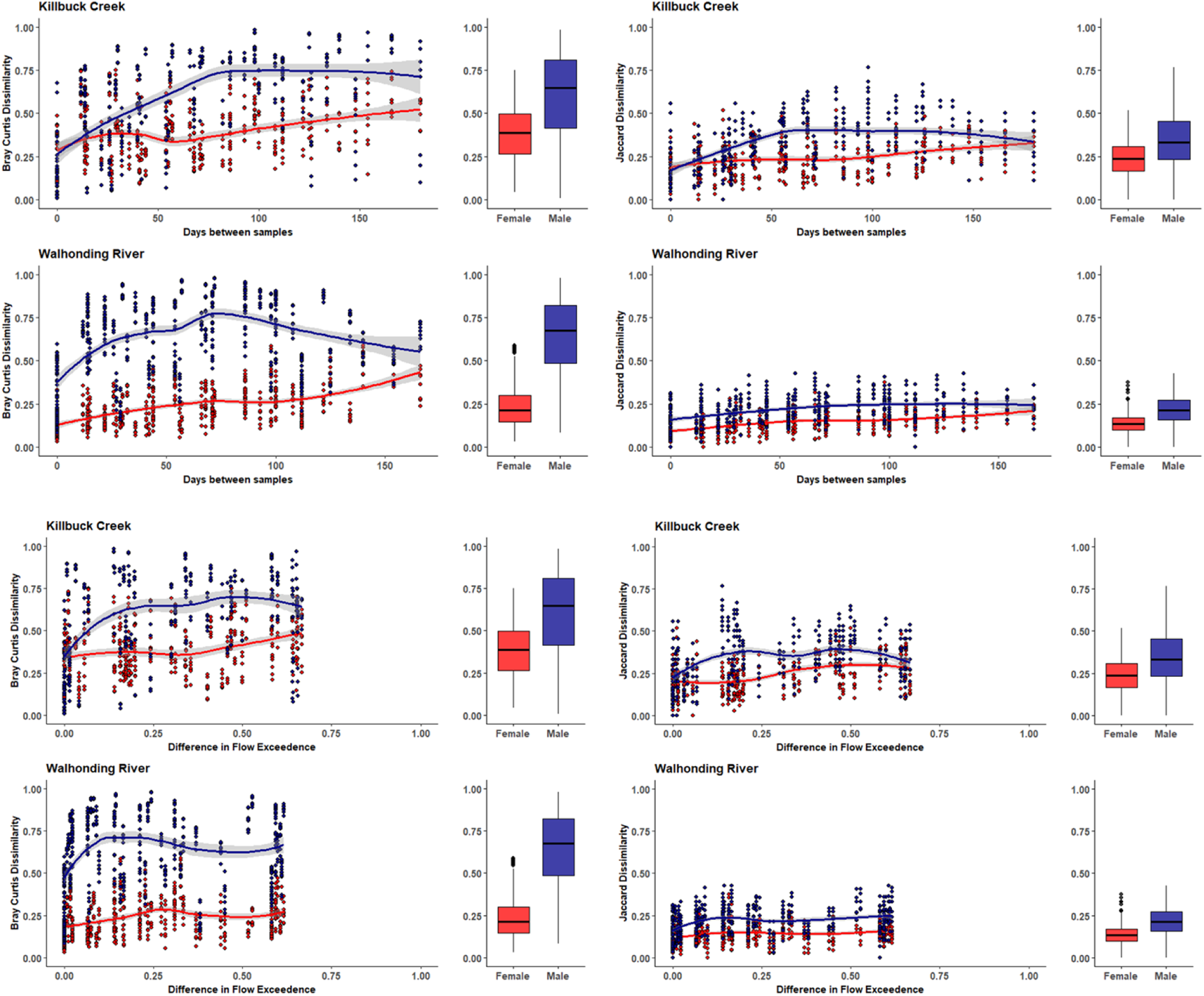
Beta diversity dissimilarity metrics calculated from the female mitotype (red) and male mitotype (blue) across the seasonal sampling events and flow exceedances within Killbuck Creek or Walhonding River. Box plots depict variance in beta diversity dissimilarities for the female and male mitotype.

**Supplementary Table 1.** Environmental DNA sample metadata for each sampling event in Killbuck Creek and Walhonding River. A measure of flow exceedance was generated by calculating the exceedance probability on each sampling data, where the exceedance probability is the total number of days in which the gage height was higher than that on the sampling date divided by the total number of days on record for that gage.

| **Sampling Site** | **Sampling Trip** | **Replicates** | **Date** | **Julian Day of**<br>**Year** | **Gage Height**<br>**(ft)** | **Flow**<br>**Exceedance** |
| --- | --- | --- | --- | --- | --- | --- |
| Killbuck Cr. | 1 | 3 | 04252024 | 116 | 9.58 | 0.675 |
| Killbuck Cr. | 2 | 3 | 05092024 | 130 | 9.02 | 0.617 |
| Killbuck Cr. | 3 | 3 | 05212024 | 142 | 9.69 | 0.686 |
| Killbuck Cr. | 4 | 3 | 06042024 | 156 | 7.59 | 0.374 |
| Killbuck Cr. | 5 | 3 | 06192024 | 171 | 6.84 | 0.216 |
| Killbuck Cr. | 6 | 3 | 07162024 | 198 | 6.69 | 0.175 |
| Killbuck Cr. | 7 | 3 | 08152024 | 228 | 6.82 | 0.207 |
| Killbuck Cr. | 8 | 3 | 08292024 | 242 | 6.17 | 0.033 |
| Killbuck Cr. | 9 | 3 | 09202024 | 264 | 6.04 | 0.019 |
| Killbuck Cr. | 10 | 3 | 10222024 | 296 | 6.20 | 0.035 |
| Walhonding R. | 2 | 3 | 05092024 | 130 | 2.66 | 0.608 |
| Walhonding R. | 3 | 3 | 05212024 | 142 | 2.67 | 0.619 |
| Walhonding R. | 4 | 3 | 06042024 | 156 | 1.47 | 0.335 |
| Walhonding R. | 5 | 3 | 06192024 | 171 | 1.13 | 0.237 |
| Walhonding R. | 6 | 3 | 07162024 | 198 | 0.92 | 0.167 |
| Walhonding R. | 7 | 3 | 08152024 | 228 | 0.70 | 0.091 |
| Walhonding R. | 8 | 9 | 08292024 | 242 | 0.43 | 0.025 |
| Walhonding R. | 9 | 3 | 09202024 | 264 | 0.20 | 0.002 |
| Walhonding R. | 10 | 3 | 10222024 | 296 | 0.24 | 0.009 |

